# Multi-omic analysis reveals the unique glycan landscape of the blood-brain barrier glycocalyx

**DOI:** 10.1101/2025.04.07.645297

**Authors:** Reid Larsen, Krzysztof Kucharz, Sidar Aydin, Mariel Kristine B. Micael, Biswa Choudhury, Mousumi Paulchakrabarti, Micael Lønstrup, Ding Chiao Lin, Markus Abeln, Anja Münster-Kühnel, Alejandro Gomez Toledo, Martin Lauritzen, Jeffrey D. Esko, Richard Daneman

## Abstract

The blood-brain barrier (BBB) glycocalyx is the dense layer of glycans and glycoconjugates that coats the luminal surface of the central nervous system (CNS) vasculature. Despite being the first point of contact between the blood and brain, not much is known about the BBB glycocalyx. Here, we performed a multi-omic investigation of the BBB glycocalyx which revealed a unique glycan landscape characterized by enrichment of sialic acid, chondroitin sulfate, and hyaluronan. We found that the BBB glycocalyx was thicker than glycocalyces in the peripheral vasculature and that hyaluronan was the major contributor to its ultrastructure. Using endothelial RNA sequencing, we found potential genetic determinants for these differences, including BBB enrichment of genes involved in sialic acid addition and peripheral enrichment of Tmem2 and Hyal2, the only known cell-surface hyaluronidases. Glycocalyx degradation and increases in vascular permeability are widely associated with inflammation. However, we found that the BBB glycocalyx remains largely unchanged in neuroinflammation during the experimental autoimmune encephalomyelitis (EAE) model of multiple sclerosis and that its degradation is not sufficient to alter BBB permeability in health. Moreover, we showed that CNS endothelial sialic acid removal delays onset of EAE, indicating that BBB glycocalyx sialic acid may contribute to the progression of neuroinflammation. These findings underscore the unique and robust nature of the BBB glycocalyx and provide targets and tools for future studies into its role in health and neuroinflammation.

---

The blood-brain barrier (BBB) describes a set of properties unique to the vasculature of the central nervous system (CNS) that tightly regulates the movement of ions, molecules, and cells between the blood and the brain^1^. The BBB is critical for maintaining brain homeostasis and its dysfunction is a hallmark of many neuroinflammatory diseases, including multiple sclerosis, epilepsy, and stroke^2^. Many BBB properties are provided by the endothelial cells (ECs) of the CNS, in conjunction with the other cells of the neurovascular unit, including mural cells, astrocytes, microglia and perivascular macrophages^3^. Several distinct components of the neurovascular unit work together to limit paracellular and transcellular transport, non-specific transcytosis, and leukocyte extravasation while simultaneously shuttling essential nutrients between the blood and the brain parenchyma^3,4^. These components include tight junctions, efflux pumps, specific nutrient transporters, transcytosis inhibitors, and low levels of leukocyte adhesion molecules^3–5^. One component of the BBB that remains poorly understood is the BBB glycocalyx, the dense glycan-rich meshwork that lines the luminal surface of the CNS vasculature^2,4,6^.

All blood vessels are lined with a glycocalyx that serves as the interface between the blood and vasculature, but the composition and properties of each glycocalyx are unique to a given vascular bed or location in the vascular tree^7–9^. The glycocalyx is composed of various glycoconjugates, particularly the long, linear, negatively charged glycans known as glycosaminoglycans (GAGs), which include heparan sulfate (HS), chondroitin sulfate (CS), and hyaluronan^10–12^. Most proteins on the luminal surface of ECs are glycosylated, with their glycans typically capped by sialic acid, a negatively charged monosaccharide that provides much of the negative charge of the cell surface^13,14^. In studies outside of the CNS, the glycocalyces of the peripheral vasculature (PV) sense blood flow, regulate blood clotting, and control the endothelial microenvironment^11,15^. Additionally, several PV glycocalyces limit vascular permeability by acting as a filtration barrier, in which their degradation increases vascular permeability^16–18^.

During inflammation, there are several proposed roles for the glycocalyx. Some studies suggest it acts as a physical barrier, whereupon its degradation in inflammation increases vascular permeability and allows leukocytes to interact with previously shielded adhesion molecules and cross the endothelium^19,20^. Conversely, the glycocalyx may also be pro-inflammatory, serving an integral role in leukocyte recruitment by enabling the formation of chemokine gradients and because several of its components, including HS and hyaluronan, serve as ligands that are essential for leukocyte interactions^7,21,22^. Therefore, the glycocalyx may have a complex role in modulating inflammation with both pro- and anti-inflammatory properties^7^.

While PV glycocalyces have been more thoroughly studied, the BBB glycocalyx is only recently gaining attention^6,8,23–25^. Recent investigations demonstrated that the BBB glycocalyx was significantly thicker than the heart and lung glycocalyces in capillaries, indicating unique characteristics of the BBB glycocalyx^26^. The BBB glycocalyx also prevented large but not small molecules from reaching the EC surface, thus serving as a molecular sieve like in the PV and acting as the first component in the BBB^24^. There is evidence for BBB glycocalyx degradation in neuroinflammatory conditions such as aging multiple sclerosis, cardiac arrest, stroke, sepsis, and cerebral malaria^25–32^. Furthermore, in stroke and cardiac arrest models, BBB glycocalyx degradation exacerbated BBB leakage and disease symptoms^27,32^. Despite these promising discoveries, the BBB glycocalyx remains a highly enigmatic and poorly understood structure, with many fundamental questions still unanswered. Which glycans are most abundant in the BBB glycocalyx, and how does molecular composition vary between glycocalyces? Which glycans contribute most to BBB structure and function? How does the BBB glycocalyx vary across the vascular tree? How does neuroinflammation affect the BBB glycocalyx?

Here, we show that the BBB glycocalyx is highly distinct from PV glycocalyces. Improving upon previous electron microscopy (EM) methods, we show that (i) the BBB glycocalyx is significantly thicker than PV glycocalyces, (ii) its structure remains relatively uniform across the vascular tree, and (iii) the presence of hyaluronan is critical for its structure. Using a novel glycomics approach, we find that the BBB glycocalyx is enriched in sialic acid, CS, and hyaluronan compared with kidney and liver glycocalyces. Using endothelial RNA sequencing, we identify potential genetic determinants for these differences, including BBB enrichment of genes involved in sialic acid addition and peripheral enrichment of *Tmem2* and *Hyal2*, the genes encoding the only known cell-surface hyaluronidases^13^. The BBB glycocalyx acts as a molecular sieve, but we find that its degradation is not sufficient to alter BBB permeability in health, so the primary barrier function of the BBB glycocalyx may be to regulate access to the CNS endothelium^24^. Contrary to previous studies, we show that neuroinflammation during the experimental autoimmune encephalomyelitis (EAE) model of multiple sclerosis leaves the BBB glycocalyx largely unchanged, even at sites of active leukocyte extravasation. Moreover, genetic removal of CNS endothelial sialic acid causes delayed onset of paralysis in EAE. This suggests that BBB glycocalyx sialic acid plays a critical role in disease progression, likely by influencing immune cell interactions at the BBB. Taken together, these findings highlight the unique structural, molecular, and functional characteristics of the BBB glycocalyx in health and neuroinflammation.

## Results

### The BBB glycocalyx is thicker than PV glycocalyces and is relatively uniform across the vascular tree

To compare BBB glycocalyx structure with that of PV glycocalyces, a modified transmission EM technique, incorporating lanthanum nitrate into the perfusion fixative, was used to enable visualization of the glycocalyx, which is absent in conventional EM^26,33–35^. The glycocalyces in the capillaries of mice were examined across several organs: the brain (cerebral cortex), muscle (hamstring), heart (left ventricle), kidney (glomerulus), and liver (sinusoids) **(Fig. 1a)**. To comprehensively assess glycocalyx structure, four key parameters were quantified for each vessel: total glycocalyx thickness, dense glycocalyx thickness, glycocalyx area (percentage of the vascular lumen), and glycocalyx coverage (percentage of the vascular surface with any glycocalyx present) **(Fig. 1b)**. The capillary BBB glycocalyx was significantly thicker and occupied more of the vessel lumen and vascular surface than the glycocalyces of all other organs that were examined. Notably, the BBB glycocalyx appeared denser closer to the endothelium, gradually thinning towards the center of the lumen. Specifically, the BBB glycocalyx had an average total thickness of 726 (±148) nm, dense thickness of 412 (±98) nm, glycocalyx area of 38 (±6.6) %, and glycocalyx coverage of 93 (±2.5) %. Glycocalyx thickness was inversely correlated with basal vascular permeability, with the fenestrated kidney glomerular and sinusoidal liver endothelia exhibiting the smallest glycocalyx and brain endothelium exhibiting the largest.

**Figure 1:**
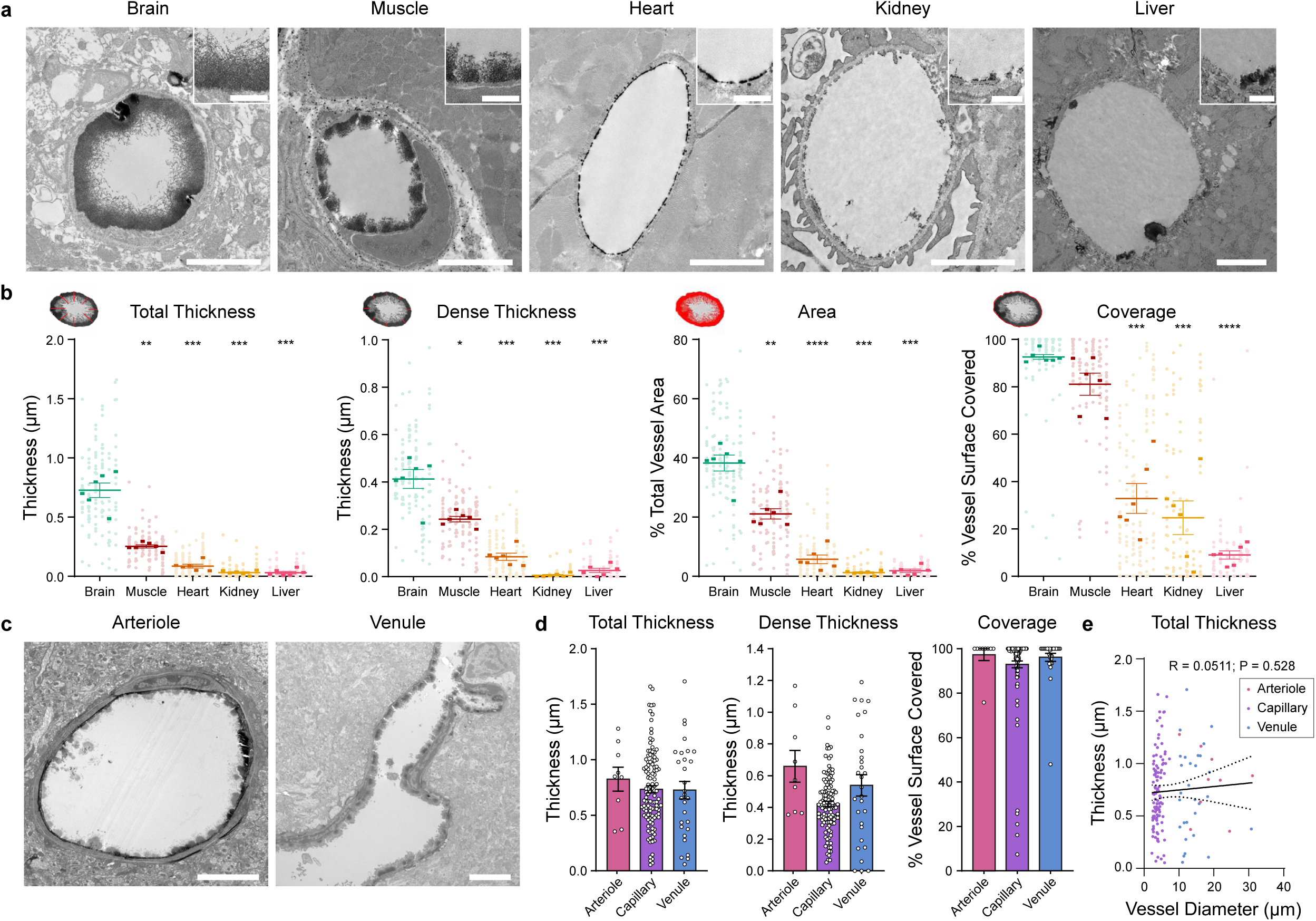
BBB glycocalyx structure is thicker than PV glycocalyces and is relatively uniform across the vascular tree. **a,** Representative lanthanum nitrate stained EM images of the glycocalyx in brain, muscle, heart, kidney, and liver capillaries (scale bar = 2 µm). Expanded view of each glycocalyx in top-right corner (scale bar = 500 nm). **b,** Quantification of glycocalyx structural parameters across organs: total glycocalyx thickness, dense glycocalyx thickness, glycocalyx area, and glycocalyx coverage. Each column represents one mouse, with each dot representing one vessel and each short bar representing the mean of a given mouse. The left three columns are female mice, and the right three columns are male (N = 6, mean ± s.e.m., *P < 0.05, **P < 0.01, ***P < 0.001, ****P < 0.0001, one-way ANOVA with multiple comparisons). **c,** Representative BBB arteriolar and venular glycocalyx images (scale bar = 5 µm) and **d** their quantifications. Each dot represents a vessel. **e,** Total thickness vs. vessel diameter of BBB arterioles, capillaries, and venules. Linear regression plotted in black with dotted 95% confidence interval (Pearson R = 0.0511, P = 0.528).

To compare BBB glycocalyx structure across the vascular tree, arterioles, capillaries, and venules were imaged by lanthanum nitrate EM. However, assessing BBB glycocalyx structure in large vessels is non-trivial due to their low density, compared to capillaries, which are by far the most abundant vessel type in the brain^1^. Moreover, in the EM specimens, the larger vessels are prone to tear due to the fragility of their large, tissue-devoid lumens. To overcome this challenge, the brains of nine additional mice were imaged, enabling the quantification of nine arterioles and 29 venules in comparison with the brain capillaries from the initial cohort **(Fig. 1c-d)**. These observations confirmed the presence of a robust glycocalyx across all segments of the vascular tree and suggested that arterioles and venules had similar structural characteristics to capillaries. The average thickness in each vessel type was independent of vascular diameter (Pearson R = 0.0511, P = 0.528) **(Fig. 1e)**. Since BBB glycocalyx thickness does not increase with increasing vascular diameter, it occupied a smaller fraction of the lumen in the large vessels compared to capillaries. Together, these data show that the BBB glycocalyx is thicker than the muscle, heart, kidney, and liver glycocalyces and that its structure remains consistent across the vascular tree in the brain.

### Endothelial transcriptomics suggest potential BBB enrichment of sialic acid and hyaluronan

The heterogeneous thickness and coverage of the glycocalyx across distinct organs suggested that the molecular composition may differ as well. RNA sequencing of brain ECs provided a global view of the glycans that might be present in the BBB glycocalyx. A set of 475 genes related to glycosylation was identified **(Supplementary File 1)** and categorized into two groups: (1) g*lycosylation genes* were defined as genes responsible for glycan synthesis, degradation, and sulfation (glycosyltransferases^36^, glycan degradation enzymes^37,38^, and (de)sulfation enzymes), and (2) *proteoglycan/mucin genes* were defined as genes encoding proteoglycan core proteins, canonical mucins, and high-confidence putative mucins^39^ (i.e. the large glycoproteins that may significantly contribute to the structure of the BBB glycocalyx). Importantly, these genes contribute to glycosylation on both the luminal (glycocalyx) and abluminal (basement membrane) side of ECs. Using previously published bulk RNA sequencing data from mice, expression levels of these genes in brain ECs were compared with those in peripheral ECs (heart, kidney, lung, and liver)^40^.

Remarkably, principal component analysis was able to separate ECs from different organs with two principal components based solely on the expression of this gene set, indicating that the endothelium of each organ may have unique glycosylation patterns **(Fig. 2a)**. Investigating expression levels of the *glycosylation genes* across organs revealed that several sialyltransferases (*St6galnac2, St8Sia4, St8Sia6, St3gal6*) and *Extl3*, a key enzyme in HS synthesis, were enriched in brain ECs compared to peripheral ECs, indicating potential enrichment of sialic acid and HS in the BBB **(Fig. 2b-c)**. The HS sulfatases *Sulf1* and *Sulf2* were highly enriched in the PV, which suggests that BBB HS may be more sulfated and therefore contribute more negative charge to the brain endothelium **(Extended Data Fig. 1)**. Peripheral ECs showed modest expression of the hyaluronan synthase genes, *Has1* and *Has3*; brain ECs showed similar *Has1* expression but lower *Has3* expression. Furthermore, peripheral ECs were enriched in several glycan degradation enzymes (*Tmem2, Hyal2, Hpse, Neu1, Mmp2, Mmp14*) **(Fig. 2b,d)**. Notably, the only known cell surface hyaluronidases *Tmem2* and *Hyal2* had approximately 20-fold and 5-fold higher expression, respectively, in peripheral ECs compared to brain ECs^13,41^. As hyaluronan is thought to be a major component of the endothelial glycocalyx^10,12^, lower brain endothelial degradation of hyaluronan could explain the increased BBB glycocalyx thickness observed by EM.

**Figure 2:**
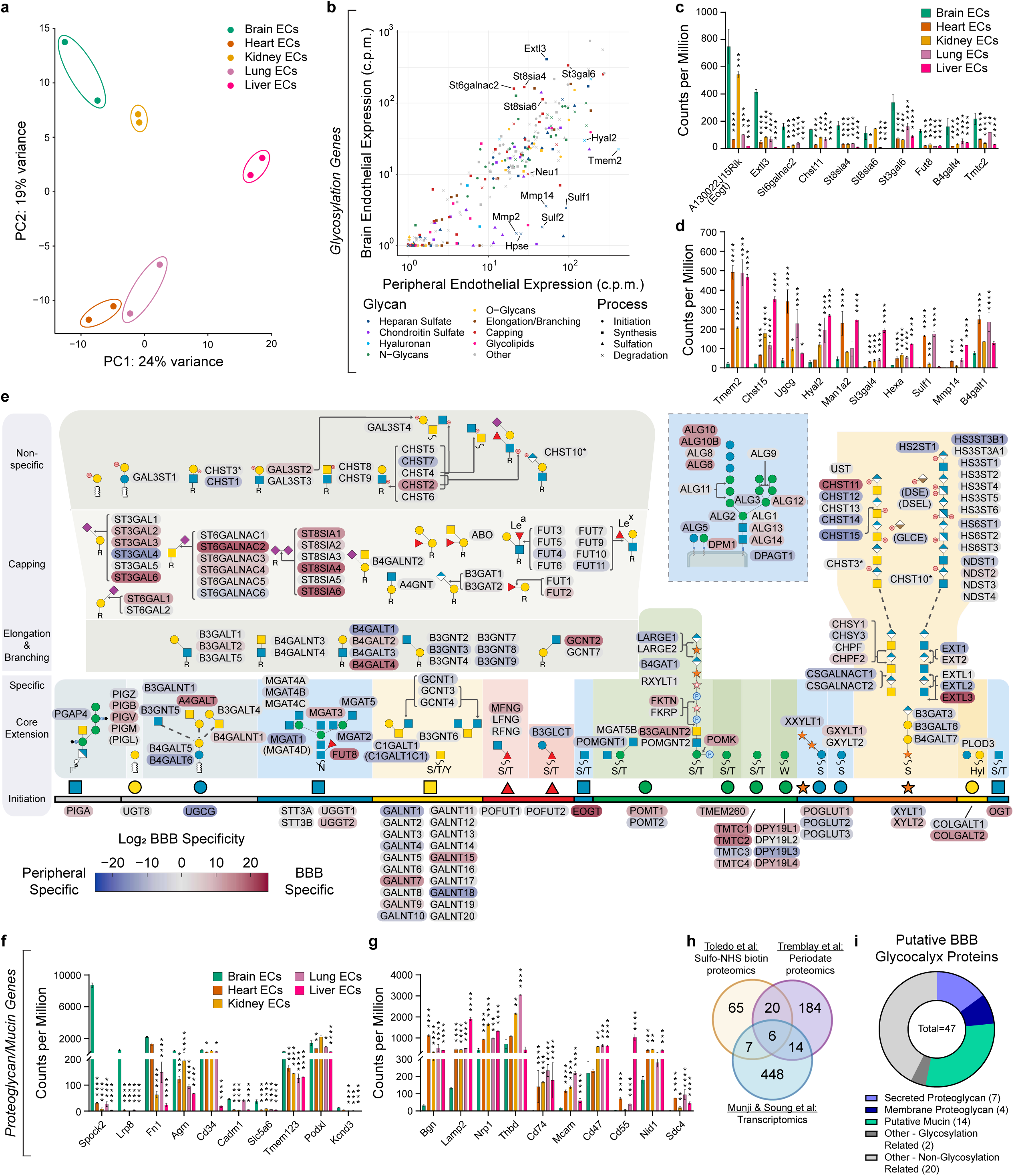
Endothelial transcriptomics suggests BBB enrichment of sialic acid and hyaluronan and the presence of several secreted proteoglycans in the BBB glycocalyx. **a,** PCA plot of ECs in brain and periphery based on expression levels of 475 glycosylation related genes. **b,** mRNA expression levels of *glycosylation genes* in brain ECs vs. peripheral ECs (average of heart, kidney, lung, and liver ECs). Genes are organized by the glycan they alter (color) and the process in which they are involved (shape). Top 10 **c** BBB and **d** peripheral specific *glycosylation genes* (N = 2, mean ± s.e.m., *FDR < 0.05, **FDR < 0.01, ***FDR < 0.001, ****FDR < 0.0001, Benjamini-Hochberg method). **e,** BBB specificity of glycosyltransferases organized by pathway with Glycopacity^42^. Red coloring indicates BBB specificity and blue coloring indicates peripheral specificity. Top 10 **f** BBB and **g** peripheral specific *proteoglycan/mucin genes* (N = 2, mean ± s.e.m., *FDR < 0.05, **FDR < 0.01, ***FDR < 0.001, ****FDR < 0.0001, Benjamini-Hochberg method). **h,** Venn diagram of glycosylation related genes and proteins identified in two luminal brain vascular proteomics datasets^44,45^. **i,** Classification of proteins found in two or three of the datasets, termed *putative BBB glycocalyx proteins.* RNA sequencing data from Munji and Suong et al^40^.

Glycopacity, a bioinformatic tool used to assess cellular glycosylation capabilities, was employed to assess the enrichment of glycan synthesis pathways^42^ **(Fig. 2e)**. This confirmed high brain EC specificity of genes involved in sialylation, with 12 sialyltransferases showing positive BBB specificity and only two showing negative BBB specificity. Though *Extl3* showed strong brain EC specificity, the genes responsible for HS polymerization (*Ext1/2*) did not. This suggests that brain ECs may be enriched in sialic acid and HS chains.

*Proteoglycan/mucin gene* expression was next examined in ECs across different organs, revealing notable BBB enrichment of *Spock2*, an extracellular matrix CS proteoglycan^43^ **(Fig. 2f-g)**. Though most canonical mucins were minimally expressed in brain ECs, some mucins, including podocalyxin (*Podxl*) and the putative mucin *Tmem123* were highly expressed. Syndecans and glypicans are membrane-bound HS proteoglycans that are generally considered important glycocalyx components^10,11^. However, only one was highly expressed (*Sdc3*) with several showing high peripheral specificity (*Sdc1/2/4, Gpc1/3/4/6*) **(Extended Data Fig. 1)**.

To further characterize the topography of proteins that may be present in the BBB glycocalyx, transcriptomic analysis was paired with two previously published studies that investigated the proteome of the luminal side of the brain vasculature^44,45^. These studies used sulfo-NHS biotin to tag proteins with exposed lysine residues^44^ or sodium periodate to tag luminal brain endothelial proteins with exposed sialic acid residues^45^ before proteomic analysis. Pairing the 475 genes derived from the transcriptomic analysis with the 98 and 224 proteins found in the luminal brain vasculature from the proteomic studies revealed that 47 proteins were identified in two or more of the studies, which were labeled *putative BBB glycocalyx proteins* **(Fig. 2h-i, Supplementary File 2)**. This group included 20 proteins not related to glycosylation, including well-known brain EC proteins, like Cdh5 and Sema7a, as well as 14 putative mucins. Notably, syndecans and glypicans were not found in the brain in either proteomics investigation, but glypicans were found in the heart, kidney, and liver^44,45^. Interestingly, seven *putative BBB glycocalyx proteins* were secreted proteoglycans, including four of the six species found in all three datasets (Hspg2, Dcn, Bgn, Col18a1). Also of note, *Spock2*, despite having higher brain EC expression than any other gene, was not identified in either proteomics dataset. This may be due to higher abundance on the abluminal side of brain ECs and the high specificity of these labeling techniques for the luminal side of the brain vasculature. In sum, these data indicate that sialic acid and hyaluronan may be enriched in the BBB and that there may be several secreted proteoglycans in the BBB glycocalyx.

### Vascular glycomics reveals enrichment of sialic acid, CS, and hyaluronan in the BBB glycocalyx

To interrogate the glycan composition of the BBB glycocalyx, a new protocol was developed based on studies investigating the vascular proteome^44,46,47^. This approach involved transcardially perfusing mice with sulfo-NHS biotin to tag the luminal vasculature **(Fig. 3a)**. From there, a tissue fraction enriched in the glycocalyx was isolated by running filtered brain, kidney, and liver homogenates over a streptavidin column, and bound biotin tagged vascular glycoproteins were eluted with trypsin. Since sulfo-NHS biotin reacts with exposed primary amines on the N-terminus or lysine residues of proteins, most vascular cell surface glycoproteins could be tagged. However, hyaluronan tagging was not anticipated as it is not covalently bound to any proteins nor possesses any primary amines itself^13^. By staining for biotin localization, preferential tagging of the vascular surface throughout the body was confirmed^44^ **(Extended Data Fig. 2a)**. Furthermore, there was a dramatic reduction in glucose from the isolated liver vasculature samples compared with pre-isolated whole liver homogenate controls where glycogen, and hence glucose, is abundant, providing further evidence of glycocalyx enrichment **(Extended Data Fig. 2b)**.

**Figure 3:**
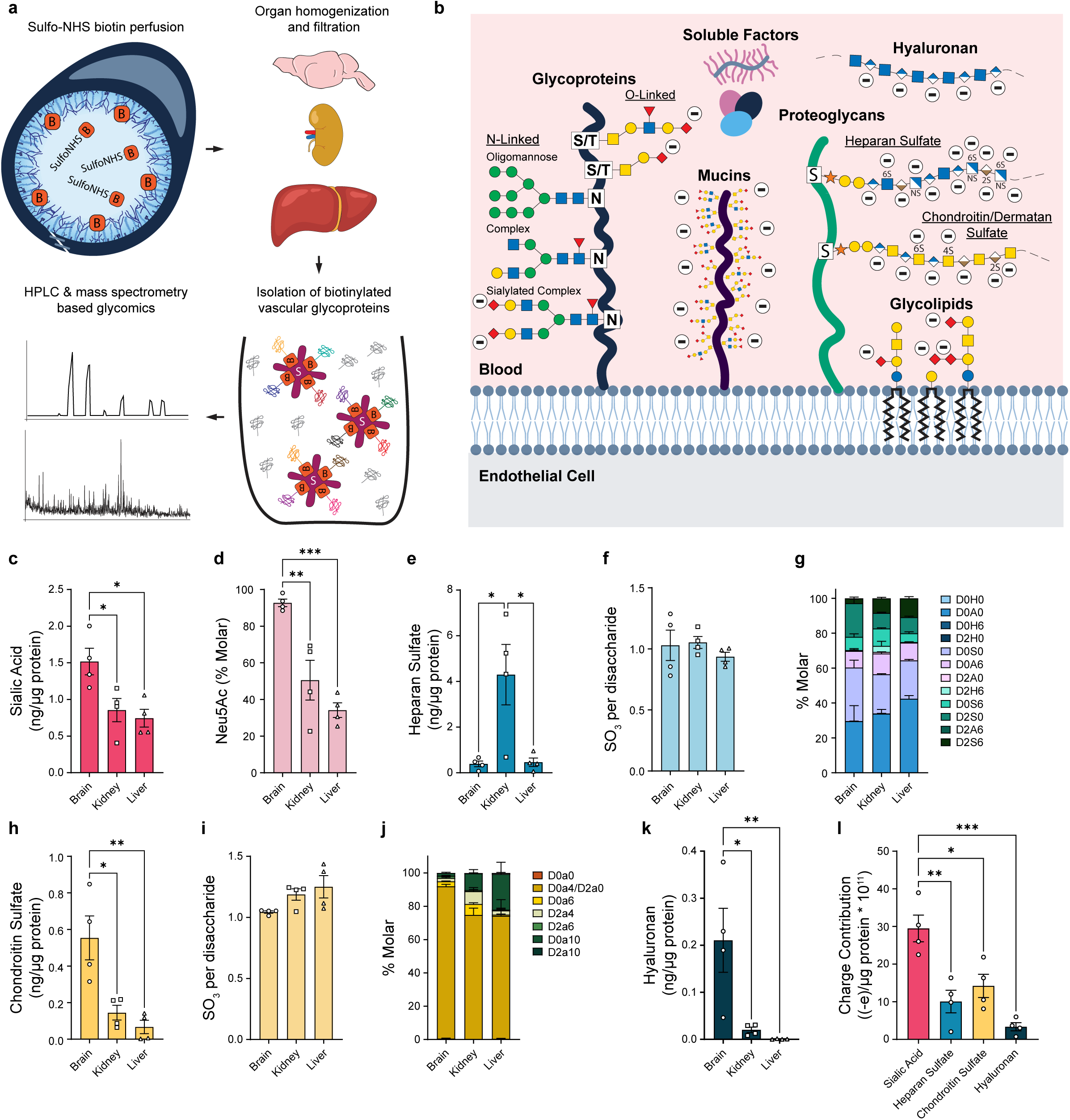
BBB glycocalyx is enriched in sialic acid, CS, and hyaluronan. **a,** Schematic of vascular glycomics protocol. Mice are transcardially perfused with sulfo-NHS biotin to covalently link biotin to glycoproteins on the luminal vasculature. The brain, kidney, and liver are isolated, homogenized and filtered before being run over a streptavidin agarose column. Biotinylated vascular glycoproteins bind to the streptavidin column while the rest of the material is washed away. Biotinylated vascular glycoproteins can then be eluted with trypsin before HPLC and mass spectrometry based glycomics. **b,** Schematic of putative glycocalyx components and where negative charge is derived. **c,** Abundance of sialic acid and **d** % Neu5Ac relative to total sialic acid. **e,h,** Abundance, **f,i** average sulfation, and **g,j,** disaccharide sulfation patterning of HS **(e-g)** and CS **(h-j)**. **k,** Abundance of hyaluronan. Glycan abundances are normalized to isolated protein following streptavidin purification. **l,** Relative charge contribution from sialic acid, HS, CS, and hyaluronan based on data in **c-k** (see methods for calculation) (N = 4, mean ± s.e.m., *P < 0.05, **P < 0.01, ***P < 0.001, one-way ANOVA with multiple comparisons).

Following glycocalyx tagging, several putative glycocalyx components, including sialic acid, HS, CS, and N-glycans, were analyzed using liquid chromatography and mass spectrometry based glycomics **(Fig. 3b)**. Normalized to biotin-tagged eluted protein, the BBB glycocalyx had almost two times more sialic acid than the kidney or liver glycocalyx, in line with predictions from our transcriptomic results **(Fig. 3c)**. While sialic acid can be found in the form of both N-acetylneuraminic acid (Neu5Ac) and N-glycolylneuraminic acid (Neu5Gc) in mice, 93% of BBB glycocalyx sialic acid was Neu5Ac on average compared with 51% in the kidney and 34% in the liver **(Fig. 3d)**. More HS was detected in the kidney vascular tagged material compared to the brain and liver, but the extent of sulfation and sulfation patterning did not differ significantly based on disaccharide analysis **(Fig. 3e-g)**. Surprisingly, CS was enriched in the BBB glycocalyx, but showed nominally less sulfation, largely driven by lower levels of 4-/6- di-sulfated N-acetyl galactosamine (GalNAc) residues (D0a10) **(Fig. 3h-j)**. N-glycan composition was also profiled in the BBB, kidney, and liver glycocalyces **(Extended Data Fig. 3a-c)**. N-glycans begin as oligomannose N-glycans, which undergo trimming to various degrees^13^. N-glycans can be elaborated into complex N-glycans, containing galactose, N-acetylglucosamine, and fucose, and are frequently capped with sialic acid to form sialylated complex N-glycans^13^. In all three organs, five oligomannose N-glycan species were identified. Eight non-sialylated complex N-glycans were detected in the BBB and kidney glycocalyces, while only two were found in the liver glycocalyx. Additionally, seven sialylated complex N-glycans were identified in the BBB glycocalyx, four in the kidney glycocalyx, and three in the liver glycocalyx. These results suggest that the liver glycocalyx may contain primarily oligomannose N-glycans and that there may be more sialylated complex N-glycan species in the BBB glycocalyx.

To assess hyaluronan levels, this experiment was repeated with the addition of 1-ethyl-3-(3-dimethylaminopropyl) carbodiimide (EDC) to the sulfo-NHS biotin solution, allowing carboxyl groups present on hyaluronan to be tagged with sulfo-NHS biotin. This enabled the capture of glycocalyx hyaluronan, which could be eluted from the streptavidin column by trypsin and hyaluronidase digestion before liquid chromatography-mass spectrometry based hyaluronan quantification. The BBB glycocalyx contained significantly more hyaluronan than the kidney and liver glycocalyces, with levels exceeding those in the kidney by more than 10-fold and hyaluronan being nearly absent in the liver glycocalyx **(Fig. 3k)**. These data are consistent with cell surface hyaluronidase gene expression of *Tmem2* (22, 207, 466 counts per million (c.p.m.) in brain, kidney, and liver ECs respectively) and *Hyal2* (29, 120, 269 c.p.m.) **(Fig. 2d)**. Based on these glycan levels, sialic acid contributed significantly more negative charge to the BBB glycocalyx than HS, CS, and hyaluronan **(Fig. 3l)**.

Histological staining of the glycans of interest provided spatial information about glycan localization in the brain (cortex, cerebellum, and hippocampus), spinal cord, muscle, heart, kidney, and liver. α2-3 linked sialic acid was prominent in the brain vasculature and parenchyma but less abundant in the larger αSMA stained vessels **(Extended Data Fig. 4a)**. In contrast, α2-6 linked sialic acid was primarily detected in the vasculature across all organs and in both capillaries and αSMA stained vessels **(Extended Data Fig. 4b)**. Both HS and mucins were generally present in the parenchymal tissue across organs with a stronger signal in the vasculature **(Extended Data Fig. 4c, 5a)**. CS was primarily found in the parenchyma with mild vascular staining in the CNS and variable staining in peripheral organs **(Extended Data Fig. 5b)**. Hyaluronan was primarily detected in the parenchyma with minimal vascular signal, except muscle and heart where there was a prominent signal between muscle fibers **(Extended Data Fig. 5c)**. Taken together, these results indicate that the glycan landscape of the BBB glycocalyx is different than that of PV glycocalyces, with BBB glycocalyx enrichment of sialic acid, CS, and hyaluronan.

### Hyaluronan is required for BBB glycocalyx structure

To identify which glycans contribute most to BBB glycocalyx structure, mice were retroorbitally injected with glycosidases to selectively degrade different glycans in the BBB glycocalyx. Sialidase, a mixture of heparinase I, II, and III, chondroitinase ABC, hyaluronidase, or mucinase (StcE) were injected to degrade sialic acid, HS, CS, hyaluronan, or mucins, respectively. Additionally, a cocktail of all five enzymes was used to degrade all five glycan classes simultaneously. Histological staining with different glycan binding proteins enabled the assessment of the efficacy of enzyme treatment **(Extended Data Fig. 6a-f)**. Sialic acid, CS, and mucins were all effectively depleted from the brain vasculature with enzymatic treatment **(Extended Data Fig. 6a**,d**,f)**. HS was detected primarily on the basement membrane of brain ECs; tamoxifen-inducible deletion of vascular exostosin like glycosyltransferase 3 *(Extl3)* did not deplete vascular HS staining, likely due to very slow basement membrane turnover in brain ECs following *Extl3* knockout **(Extended Data Fig. 7a-b)**. However, HS was depleted when primary brain ECs were cultured *in vitro*, following digestion with collagenase/dispase, which degraded the basement membrane **(Extended Data Fig. 7c-e)**. This observation was confirmed by high magnification thin section immunohistochemistry, which revealed HS localization primarily on the abluminal side of brain ECs **(Extended Data Fig. 7f)**. Thus, as expected, the BBB prevented heparinase-dependent removal of HS in the BBB basement membrane **(Extended Data Fig. 6b)**. However, heparinase treatment degraded basement membrane HS in the kidney, which does not have a BBB to prevent access to basement membrane HS **(Extended Data Fig. 6c)**. Therefore, any HS present in the BBB glycocalyx was likely degraded following heparinase injection. Interestingly, there was diffuse hyaluronan staining throughout the tissue with or without hyaluronidase injection **(Extended Data Fig. 6e)**.

After glycan degradation, BBB glycocalyx structure was assessed by EM with lanthanum nitrate staining **(Fig. 4a)**. Compared to uninjected controls (670 ± 77 nm total thickness), injection of sialidase (582 ± 172 nm) and chondroitinase (639 ± 33 nm) did not significantly alter BBB glycocalyx thickness **(Fig. 4b)**. Injection of heparinase (461 ± 256 nm) and mucinase (496 ± 105 nm) caused a modest though non-statistically significant reduction in BBB glycocalyx thickness. Remarkably, injection of hyaluronidase significantly reduced the thickness of the BBB glycocalyx (183 ± 162 nm), removing much of its bushy structure and leaving a thin glycocalyx layer at the surface of the ECs. Injection of a cocktail of all the enzymes showed a similar effect (205 ± 100 nm); however, much of the EC surface showed no presence of glycocalyx (57 ± 11% average coverage). These data suggest that the presence of hyaluronan is critical to BBB glycocalyx structure, particularly in the non-dense portion furthest from the brain endothelium.

**Figure 4:**
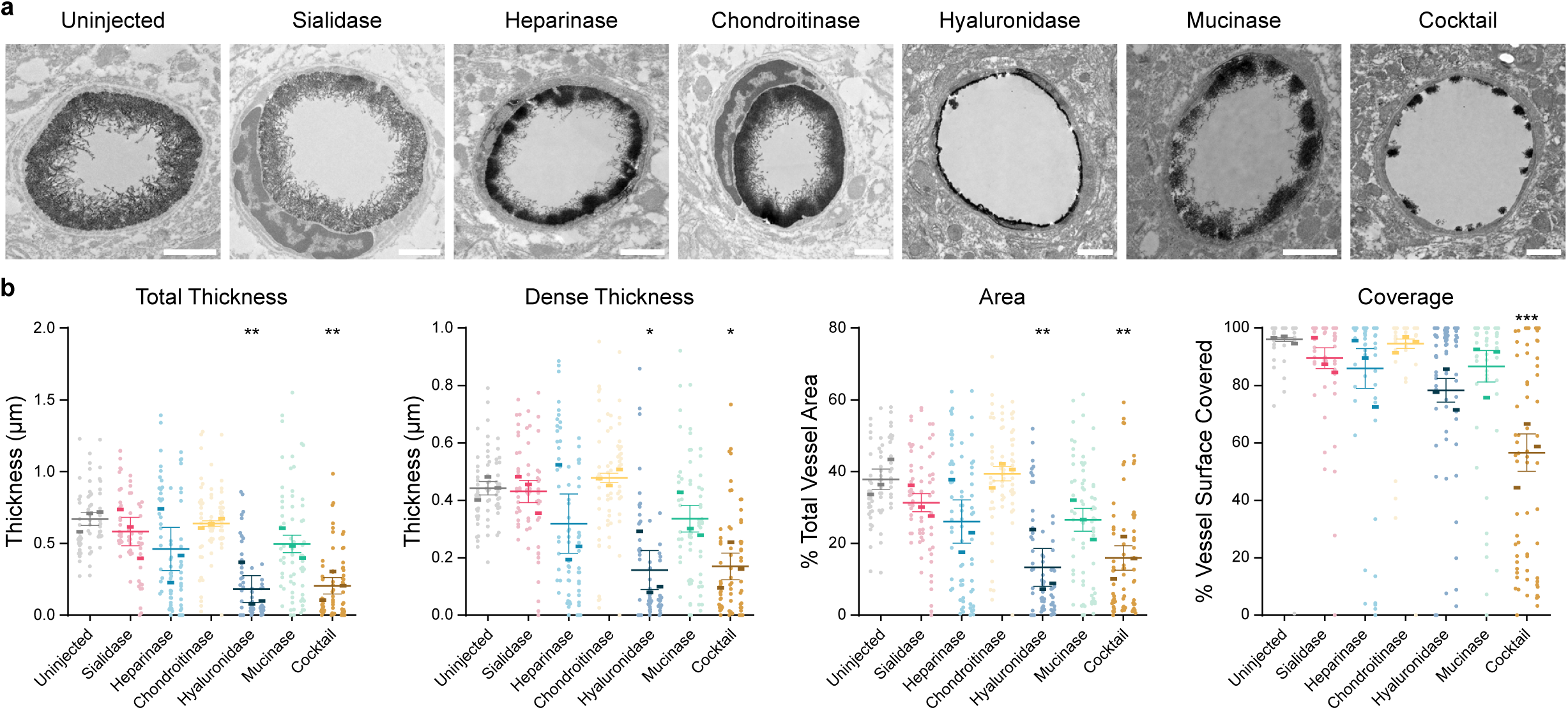
Hyaluronan is necessary for maintenance of BBB glycocalyx structure. **a,** Representative lanthanum nitrate stained EM images of BBB glycocalyx in cerebral cortex capillaries following retroorbital injection of the indicated enzymes (scale bar = 1 µm). **b,** Quantification of BBB glycocalyx parameters after enzyme injection (N = 3, mean ± s.e.m., *P < 0.05, **P < 0.01, ***P < 0.001, one-way ANOVA with multiple comparisons).

### BBB glycocalyx degradation does not increase BBB permeability in health

To determine whether degradation of the BBB glycocalyx alters BBB permeability, glycans were selectively removed from the BBB glycocalyx by retroorbital injection of glycosidases, as before. Mice were then injected with sodium fluorescein (NaFl, 332 Da), perfused with PBS, and NaFl fluorescence was measured with a spectrophotometer in brain homogenates and blood plasma. This yielded a ratio of brain:blood fluorescence intensity, which was used to quantify BBB permeability. BBB permeability of NaFl did not increase after injection of any of the enzymes **(Fig. 5a)**. Next, the BBB was assessed with *in vivo* two-photon microscopy, a more sensitive technique that can assess BBB permeability in the intact, living brain^48^. To prepare for imaging, mice were ventilated, arterial and venous catheters were inserted for delivery of tracers and anesthesia, respectively, and a cranial window was made over the somatosensory (barrel) cortex of the brain **(Fig. 5b)**. The BBB glycocalyx was then manipulated by retroorbital injection of PBS (control, intact glycocalyx), enzyme cocktail (largely degraded glycocalyx), or sialidase (removed sialic acid), given the emerging evidence of the role of sialic acid in vascular permeability^16^ and its high abundance and charge contribution in the BBB glycocalyx **(Fig. 3c,l)**.

**Figure 5:**
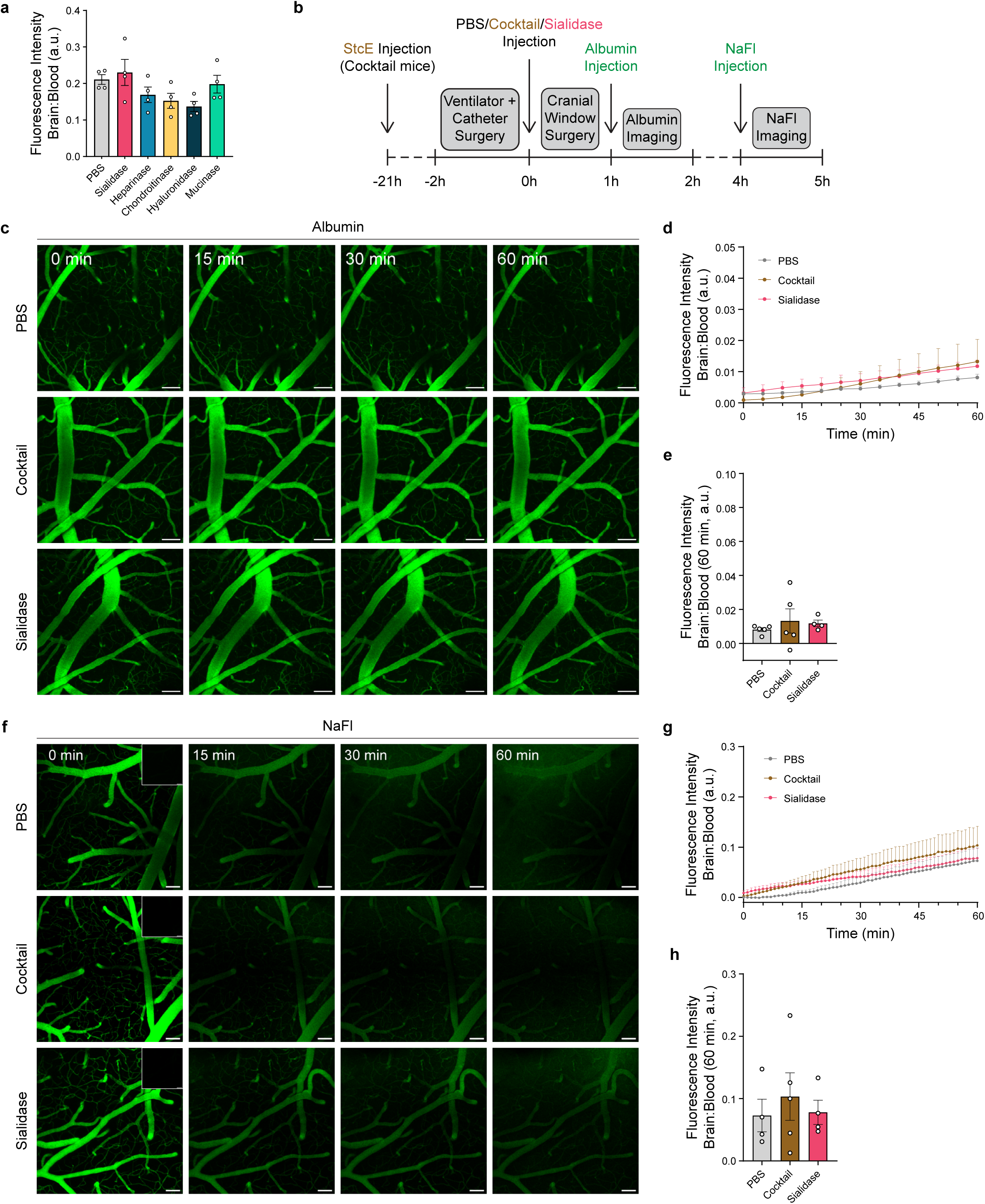
Degradation of the BBB glycocalyx does not alter BBB permeability in healthy mice. **a,** Relative fluorescence intensities of NaFl in brain homogenate vs. blood plasma in mice treated with PBS or the specified enzymes as measured by spectrophotometer (N = 4, mean ± s.e.m., one-way ANOVA with multiple comparisons). **b,** Timeline of procedure for two-photon imaging to assess BBB leakage of albumin and NaFl. **c,** Representative images from time-lapse imaging immediately after albumin injection (0 min), and 15, 30, and 60 minutes thereafter (scale bar = 100 µm). **d,** Ratio of albumin fluorescence intensity (brain parenchyma/brain vasculature) over time. **e,** Ratio of fluorescence intensity at 60 minutes after albumin injection (N = 4-5, mean ± s.e.m., one-way ANOVA with multiple comparisons). **f,** Representative images from time lapse imaging immediately after NaFl injection (0 min), and 15, 30, and 60 minutes thereafter (scale bar = 100 µm). (Top right of 0 minutes) Image of the same ROI immediately before NaFl injection. **g,** Ratio of NaFl fluorescence intensity (brain parenchyma/brain vasculature) over time. **h,** Ratio of fluorescence intensity 60 minutes after NaFl injection (N = 4-5, mean ± s.e.m., one-way ANOVA with multiple comparisons).

To measure BBB permeability, Alexa Fluor 488 conjugated bovine serum albumin, a ∼70 kDa negatively charged protein, was injected and the ratio of parenchymal fluorescence intensity to plasma fluorescence intensity in the top 100 µm of the brain was measured. Injection of the enzyme cocktail or sialidase did not lead to an increase of albumin transport across the BBB, with parenchymal levels of albumin signal remaining low in all conditions **(Fig. 5c-e, Supplementary File 3)**. Subsequently, in the same set of mice, BBB permeability to NaFl was investigated. Because the fluorescence of NaFl was substantially higher than the Alexa Fluor 488 conjugated albumin, the microscope’s recording sensitivity was reduced, effectively making the NaFl imaging insensitive to the previously injected Alexa Fluor 488 conjugated albumin **(Fig. 5f, top right of 0 min)**. Immediately following NaFl injection, strong fluorescence was observed in the brain vasculature **(Fig. 5f, 0 min)**. Over the next 60 minutes, there was moderate parenchymal NaFl accumulation in all three conditions, with a small trend towards an increase in permeability in the cocktail-injected group **(Fig. 5f-h, Supplementary File 4)**. Although BBB glycocalyx degradation has been shown to increase access to the brain endothelium^24^, these results indicate that degradation of the BBB glycocalyx does not significantly increase BBB permeability in healthy mice.

### BBB glycocalyx structure and composition are largely unchanged in EAE

Recent reports have suggested that the BBB glycocalyx may be shed in neuroinflammatory conditions^25,27,29,49,50^. To assess the BBB glycocalyx in neuroinflammation, mice were injected with pertussis toxin and myelin oligodendrocyte glycoprotein (MOG_35-55_) in Complete Freund’s Adjuvant to induce EAE, a well-established model of multiple sclerosis with robust neuroinflammation, BBB dysfunction, and leukocyte extravasation^51,52^. First, BBB glycocalyx structure was analyzed using EM with lanthanum nitrate staining at the onset, peak, and chronic timepoints of the disease in the white matter of the lumbar spinal cord, where EAE myelin lesions are most prominent^53,54^. As expected, robust myelin damage was observed in the EM specimens **(Extended Data Fig. 8a-d)**. Surprisingly, there was no significant degradation of the BBB glycocalyx at any timepoint **(Fig. 6a)**. Rather, BBB glycocalyx structure was remarkably preserved, even in venules with evident neuroinflammation where leukocytes were actively extravasating **(Fig 6b-c, Extended Data Fig. 8e-l)**. Indeed, venular BBB glycocalyx thickness and coverage remained consistent whether leukocytes were observed in the vessel or not **(Extended Data Fig. 8m-t)**. In contrast to images of healthy mice in which leukocytes are removed during perfusion, leukocytes were abundant in the vessels of EAE mice and often appeared in contact with the endothelium. This was likely due to EC-leukocyte interactions during various stages of leukocyte extravasation, which is common in EAE. Interestingly, there was a robust glycocalyx present on most leukocytes observed by EM **(Extended Data Fig. 8e-j)**.

**Figure 6:**
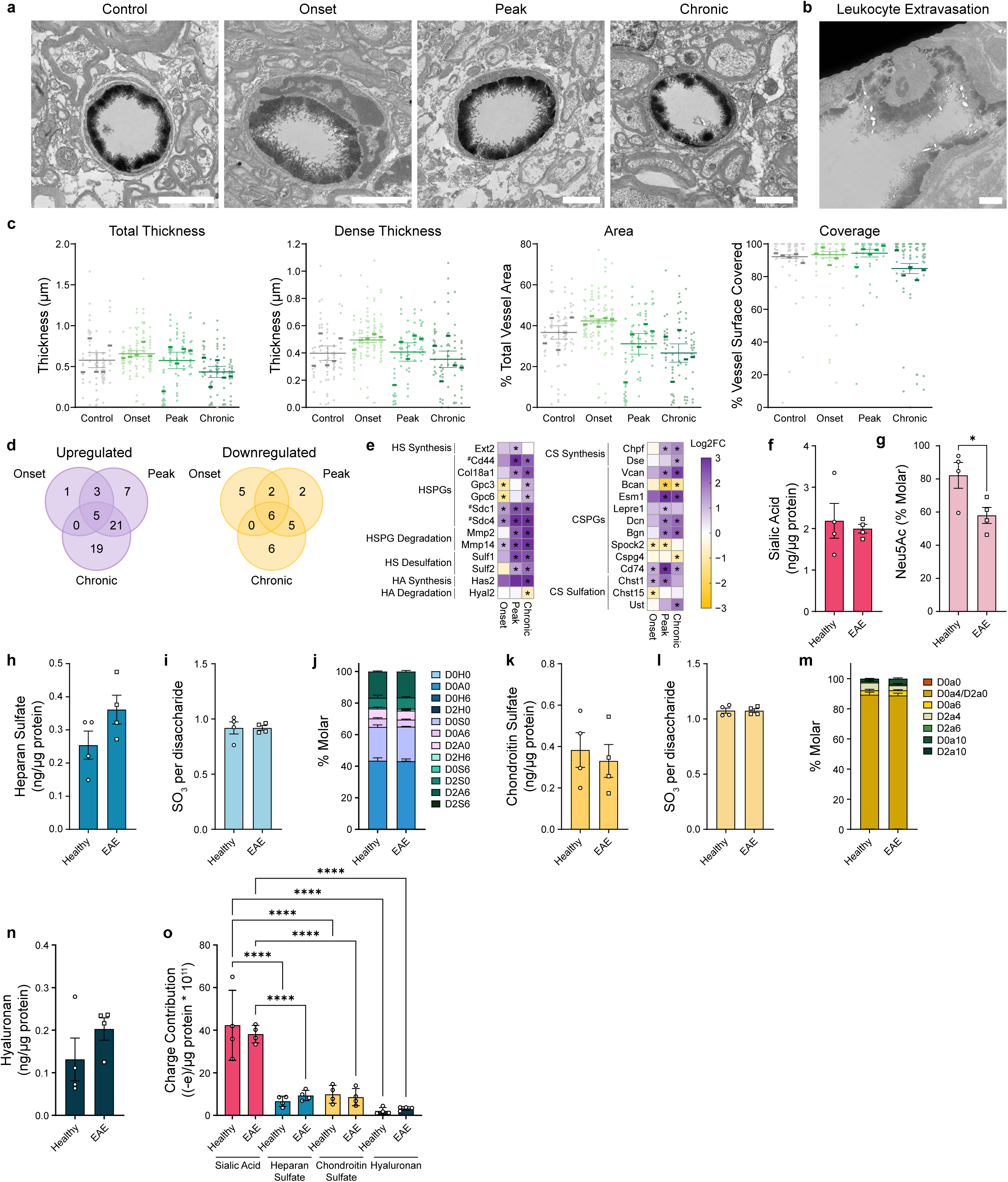
BBB glycocalyx remains largely intact during EAE even at sites of active leukocyte extravasation. **a,** Representative lanthanum nitrate stained EM images of BBB glycocalyx in white matter capillaries in lumbar spinal cord from uninjected healthy control mice, and EAE mice at onset, peak, and chronic timepoints of disease (scale bar = 2 µm). **b,** Representative image of leukocyte extravasation in EAE during peak timepoint (scale bar = 2 µm). **c,** Quantification of BBB glycocalyx parameters across timepoints of EAE (N = 5, mean ± s.e.m., one-way ANOVA with multiple comparisons). **d,** Venn diagram of glycosylation related genes that are significantly upregulated and downregulated at each timepoint in EAE. **e,** Relative change in expression of significantly altered proteoglycan genes in EAE at each timepoint. * indicates a significant up/downregulation and # indicates HS proteoglycans that also contain CS chains. Original data for **d-e** from Munji and Suong et al^40^. **f-m,** Glycomic analysis of BBB glycocalyx in the spinal cord of healthy and peak EAE mice including **f** abundance of sialic acid, **g** % Neu5Ac, **h** HS abundance, **i** sulfation, and **j** sulfation pattern, **k** CS abundance, **l** sulfation, and **m** sulfation pattern. **n,** Hyaluronan abundance (N = 4, mean ± s.e.m., *P < 0.05, two-sided t-test). **o,** Relative charge contribution from sialic acid, HS, CS, and hyaluronan based on data in **f-n** (N = 4, ****P < 0.0001, two-way ANOVA).

These unexpected findings prompted an investigation into the molecular changes to the BBB glycocalyx in EAE. First, expression of the 475 glycosylation related genes was assessed in spinal cord ECs in EAE using a previously published dataset^40^. Across all three timepoints, 56 genes were upregulated and 26 were downregulated **(Fig. 6d)**. Notably, many genes encoding proteoglycan core proteins (*Cd44, Col18a1, Dcn, Vcan*) and proteoglycan/glycoprotein degradation enzymes (*Mmp2, Mmp14*) were upregulated in EAE, leading to speculation that the BBB glycocalyx may undergo rapid turnover in EAE, rather than being purely broken down. Indeed, among the significantly altered proteoglycan-related genes, most of them were upregulated, suggesting increased synthesis of proteoglycans and GAGs in EAE **(Fig. 6e)**.

To further investigate BBB glycocalyx composition in EAE, glycomic profiles of the spinal cord glycocalyces in peak EAE and healthy mice were compared, as before. Biotin staining remained mostly vascular, though there was minor leakage in EAE mice **(Extended Data Fig. 2c)**. Sialic acid levels remained constant, but EAE mice showed a significantly lower percentage of the Neu5Ac form of sialic acid **(Fig. 6f-g)**. In line with EM and transcriptomic results, there was a nominal but non-statistically significant increase in HS levels but similar sulfation patterning in EAE mice **(Fig. 6h-j)**. Additionally, CS levels and sulfation patterning between groups remained similar **(Fig. 6k-m)**. Hyaluronan levels showed a trend towards a small increase in EAE, again in line with EM and transcriptomic results **(Fig. 6n)**. Like in the brain, sialic acid in the spinal cord contributed more negative charge to the BBB glycocalyx than HS, CS, and hyaluronan, but the charge contributions of each glycan did not change significantly in EAE **(Fig. 6o)**. Lastly, no major changes to N-glycan patterning between conditions were observed **(Extended data Fig. 9a-b)**. In conclusion, the structure and molecular composition of the BBB glycocalyx remain largely unchanged in EAE.

### CNS endothelial sialic acid depletion delays EAE onset

Sialic acid is implicated in leukocyte extravasation, with evidence for its involvement in the leukocyte rolling, firm adhesion, and chemokine mediated EC activation^14,55^. This raised the hypothesis that removal of sialic acid from the BBB glycocalyx may disrupt leukocyte extravasation and affect the progression of neuroinflammation. To test this, CNS endothelial sialic acid knockout mice were generated by targeting cytidine monophosphate N-acetylneuraminic acid synthase (Cmas), a key enzyme in sialic acid synthesis^56,57^. EAE was then induced in CNS endothelial sialic acid knockout mice (Cmas^f/f^; Slco1c1-Cre) and their littermate controls (Cmas^f/f^). Remarkably, CNS endothelial sialic acid knockout mice showed a delayed onset of paralysis **(Fig. 7a-b)**. Average time to disease onset and to hind limb paralysis were both significantly increased **(Fig. 7c-d)**. On day 13, all littermate control mice showed partial or full hind limb paralysis compared to just 4/9 in the knockout group, with another 4/9 showing tail-only paralysis and one with no paralysis **(Fig. 7e)**. Though disease onset was delayed, mice in both groups showed similar levels of paralysis by day 17, with similar maximum disease scores and total disease burden **(Fig. 7f-g)**. In summary, these data show that CNS endothelial sialic acid depletion delays EAE onset.

**Figure 7:**
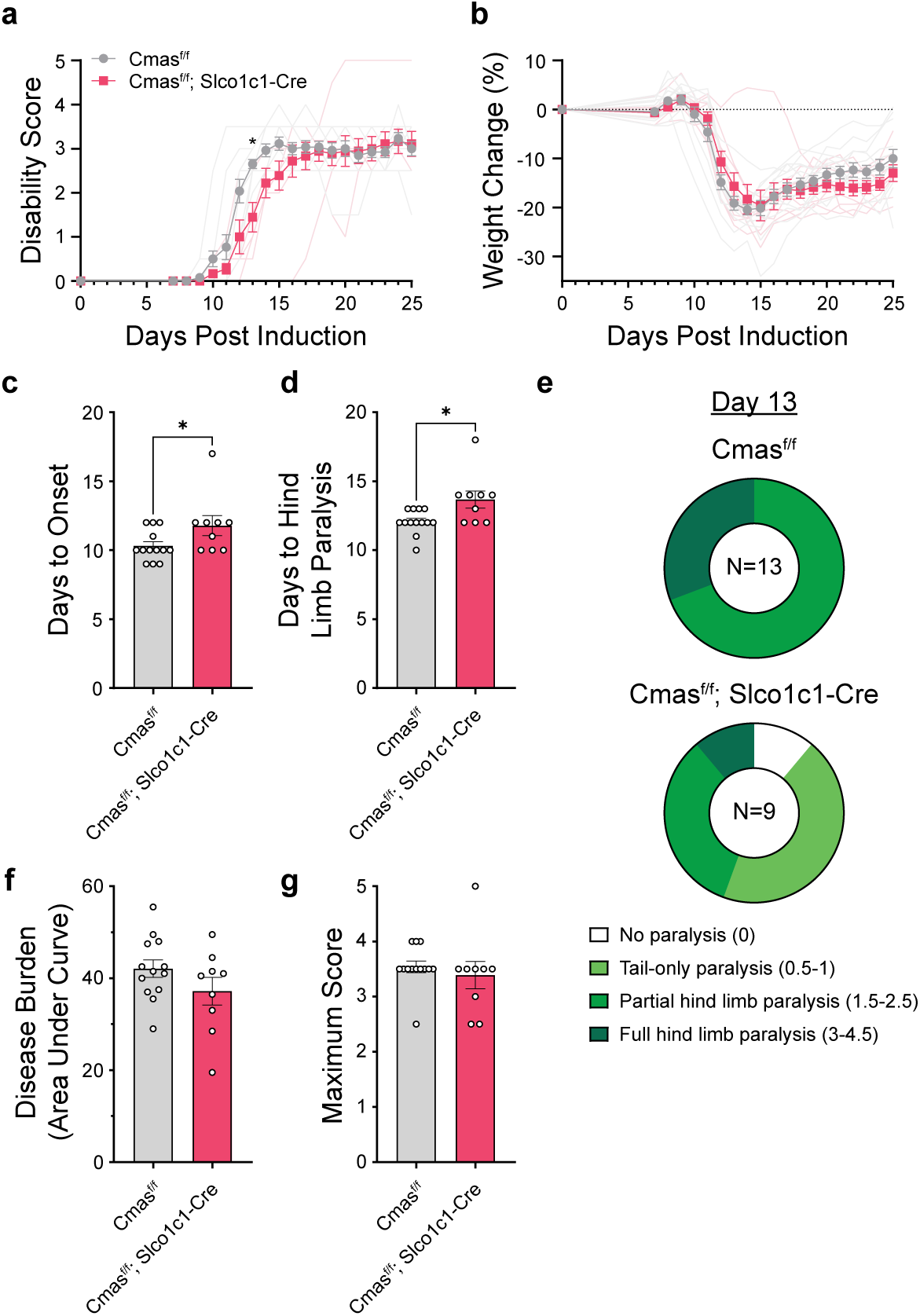
CNS endothelial sialic acid removal delays EAE onset. **a,** Disability score and **b** relative weight change in CNS endothelial sialic acid knockout (Cmas^f/f^, Slco1c1-Cre) and littermate control (Cmas^f/f^) mice after EAE induction (N = 9-13, mean ± s.e.m., *P-adj. < 0.05, multiple two-sided t-tests). **c,** Days to onset of disease (day of first scoring or 1 g weight loss over 2-day period) and **d** days to hind limb paralysis (disability score ≥ 1.5) (N = 9-13, mean ± s.e.m., *P < 0.05, two-sided t-test). **e,** Disability scores on day 13. **f,** Total disease burden (sum of daily disability scores during entire experiment) and **g** maximum disability score (N = 9-13, mean ± s.e.m., two-sided t-test).

## Discussion

The BBB glycocalyx is an understudied but potentially critical component of the BBB^2,6^. Understanding the unique features of the BBB glycocalyx could provide novel insights into the mechanisms underlying BBB dysfunction and offer new therapeutic avenues to modulate neuroinflammation. Recent technological advancements in glycobiology now enable a more detailed exploration of the structural and molecular properties of the BBB glycocalyx^34,36,44^. In this study, we leveraged and refined these approaches to gain deeper insights into the BBB glycocalyx in both healthy and neuroinflammatory states. We found that the BBB glycocalyx was distinct from PV glycocalyces, with a thicker structure and enrichment of sialic acid, CS, and hyaluronan **(Fig. 8)**. Additionally, we determined that hyaluronan is the major contributor to its structure. We showed that BBB glycocalyx degradation does not alter BBB barrier function in a healthy brain and that it remains largely unchanged in the EAE model of multiple sclerosis. However, mice that lack CNS endothelial sialic acid showed delayed paralysis in EAE, potentially driven by impaired leukocyte extravasation which depends on sialylated glycan recognition by selectins expressed on ECs and leukocytes.

**Figure 8:**
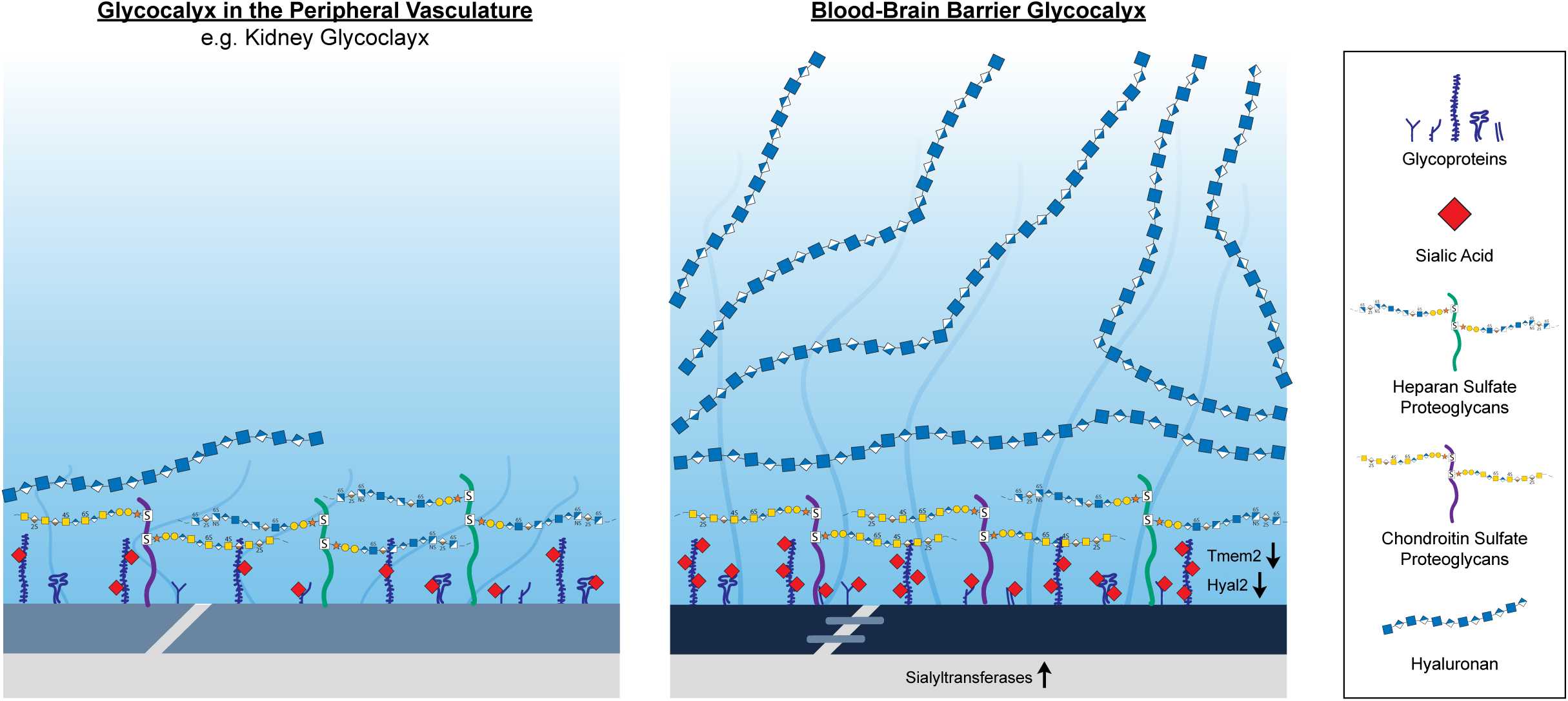
Proposed model of the BBB glycocalyx. Here we present a schematic representation of the proposed structure and molecular composition of the BBB glycocalyx, highlighting specific glycans and glycoconjugates that are the focus of this study. The BBB glycocalyx is thicker than glycocalyces in the PV and is enriched in sialic acid, chondroitin sulfate, and hyaluronan. The bulk of BBB glycocalyx ultrastructure is derived from the increased abundance of hyaluronan, particularly further from the endothelial surface. The distinct glycan landscape of the BBB glycocalyx likely comes from differential expression of several glycosylation related genes relative to the PV including lower expression of the cell surface hyaluronidases Tmem2 and Hyal2 and increased expression of sialyltransferases.

Accurate structural assessment of the endothelial glycocalyx is notoriously difficult^58–60^. Measuring glycocalyx structure in live organisms requires inference of glycocalyx location based on the exclusion of dyes or blood cells or the use of fluorescent lectins, which only bind to a subset of the glycans present in the glycocalyx^6^. Glycan specific EM provides unmatched resolution and non-specifically stains the glycocalyx based on charge^34^. However, its intensive preparation method requires a strong dehydration process, which is known to compress the glycocalyx and lead to underestimation of glycocalyx thickness^59,61^. To overcome these limitations, we modified previously published EM preparations^26,33,34^, specifically increasing the perfusion rate to more closely match the rate of a living mouse, thus initiating rapid fixation and limiting the shear stress that accompanies a rapid decrease in perfusion rate and blood pressure during transcardial perfusion. In doing so, we measured an average BBB glycocalyx thickness of 726 (±148) nm, greater than any other BBB glycocalyx EM study and more in line with estimations of glycocalyx thickness in live animals^12,60,62^. We speculate that this is due to preservation of the layer of secreted GAGs and soluble factors that forms much of the BBB glycocalyx^7^. We were also able to make the first ultrastructural comparison of BBB glycocalyx thickness across the vascular tree and found that its structure remained consistent in arterioles, capillaries, and venules.

The molecular composition of the BBB glycocalyx remains poorly understood, largely due to technical challenges in exclusively probing the native glycocalyx. Traditional methods, such as antibody or lectin staining, and most *in vitro* approaches often struggle to distinguish between the abluminal and luminal surfaces of ECs. Similarly, measurements of shed glycocalyx components in the bloodstream provide only indirect information, as these molecules can originate from other vascular beds or blood cells in the body. In our study, we addressed these limitations by employing complementary approaches to specifically investigate the BBB glycocalyx on the luminal surface of CNS ECs. In addition to conventional RNA sequencing and glycan staining, we developed a biotin-based glycomic approach to isolate and characterize luminal endothelial glycans. Further, we selectively degraded specific glycans to assess their contribution to the ultrastructure of the BBB glycocalyx. These methodologies allowed us to achieve a more precise understanding of the molecular composition of the BBB glycocalyx and highlighted sialic acid, CS, and hyaluronan as enriched components of the BBB glycocalyx. We further showed that the presence of hyaluronan is critical to BBB glycocalyx structure and suggest that lower BBB expression levels of the only known cell surface hyaluronidases, *Tmem2* and *Hyal2*, may underlie the higher BBB glycocalyx thickness and enrichment of hyaluronan. By comparing our endothelial transcriptomic analysis with two luminal brain vascular proteomics datasets, we identified seven secreted proteoglycans in two or more of the datasets and four in all the datasets. If much of the BBB glycocalyx is made of hyaluronan and secreted proteoglycans that are not covalently bound to the EC surface, this could explain the fragility that is often associated with the glycocalyx^10,11^. Based on these findings, we speculate that hyaluronan may make up the bulk of the BBB glycocalyx ultrastructure, with other secreted proteoglycans enmeshed in its structure and glycoproteins situated closer to the EC surface.

The glycocalyx is traditionally thought of as a molecular sieve regulating access to the endothelium^10,11^. In the periphery, this filtration helps limit vascular permeability, and thus disruption of several PV glycocalyces causes increased vascular permeability^12,15^. The BBB glycocalyx is also known to provide this sieve function, limiting access of large molecules to the EC surface^24^. However, its role in regulating permeability across the entire neurovascular unit was previously unknown. We showed that degradation of the BBB glycocalyx does not cause a significant increase in BBB permeability in healthy mice. Similarly, BBB permeability to Evans Blue did not increase after LPS-induced BBB glycocalyx degradation^26^. We suspect that this is because other components of the BBB remained intact and were sufficient to limit the permeability of the injected tracers. The BBB glycocalyx may play a more prominent role in regulating vascular permeability during neuroinflammation accompanied by BBB dysfunction, as has been shown in models of cardiac arrest and stroke^27,32^.

There is mounting evidence of the importance of the glycocalyx in inflammation throughout the body^11,18,63,64^. Several groups have found that PV glycocalyces are degraded in inflammatory conditions such as sepsis, trauma, diabetes, and others^11^. Glycocalyx shedding in inflammation is proposed to regulate immune cell access to the endothelium; under healthy conditions, the glycocalyx physically blocks this interaction^12,20^. On the contrary, some studies have found that glycocalyx shedding on leukocytes, rather than the endothelium, is what drives leukocyte adhesion^65^. This is less well studied in the CNS during neuroinflammation. Loss of the HS proteoglycan, syndecan 1, on leukocytes drives extravasation in the retinal vasculature^66^. In another study, the BBB glycocalyx, but not the heart and lung glycocalyces, remained partially intact following intense inflammation caused by high-dose LPS administration^26^. In contrast, several other groups found that the BBB glycocalyx was degraded in neuroinflammation during cerebral edema, stroke, cardiac arrest, and epilepsy as measured by EM and serum levels of specific glycans and glycoconjugates^27,32,50,67^. However, EM preparations can disrupt the integrity of the glycocalyx and many components are not covalently bound to the EC surface^11,59,68^. Therefore, it can be difficult to get an accurate and consistent picture of the glycocalyx with EM, particularly in disease, as this often leads to loss of vessel integrity and collapse of the vessels. Moreover, elevated serum glycan levels may not be a result of BBB glycocalyx degradation but rather faster BBB glycocalyx turnover, glycocalyx shedding on blood cells, or effects elsewhere in the body. With our modified EM preparation, we have been able to limit vessel collapse, even in sick mice and in inflamed vessels with leukocytes in the process of extravasation. With this maintained vessel integrity, we showed that the BBB glycocalyx structure remains largely unchanged in EAE. Furthermore, our glycomics results showed that at the luminal EC surface, GAG levels are not dramatically reduced, which suggests that the BBB glycocalyx composition remains relatively unchanged in EAE, even if it is simultaneously being turned over. These experiments support the idea that, at least in the spinal cord during EAE, the BBB glycocalyx may serve to facilitate inflammation, in contrast to the idea that its degradation is required for leukocyte extravasation. Indeed, leukocytes could contact the endothelium with an intact BBB glycocalyx by simply compressing it locally at the site of adhesion. Furthermore, CNS endothelial sialic acid removal was sufficient to delay EAE onset, potentially by impairing leukocyte-EC interactions. These findings challenge the notion that the BBB glycocalyx is merely degraded in neuroinflammation. Instead, it may undergo remodeling and play a more nuanced role, incorporating both pro-inflammatory components, such as sialic acid, and anti-inflammatory components, such as hyaluronan.

There are limitations to our investigation. Though our EM experiments provide a robust quantitative assay to measure BBB glycocalyx structure, it should not be assumed that it provides absolute quantification of BBB glycocalyx thickness. The EM preparation requires intense dehydration steps, which are known to compress the BBB glycocalyx^59,61^. Next, our endothelial transcriptomic investigation, though illuminating, does not provide spatial information (luminal/abluminal, vascular tree location, organ region). Furthermore, it is impossible to determine glycosylation patterns based solely on transcriptomic data, as glycans are not directly encoded in the genome and require a complex network of enzymes and substrates to be synthesized. Though our glycomic investigation preferentially targets the luminal vasculature, it does not provide spatial information across the vascular tree or location within a given organ. Additionally, though our glycomic isolation approach showed largely vascular tagging, we cannot exclude the possibility that the biotin may also have tagged some basement membrane glycans, particularly in the kidney and liver. As such, we tried to assess glycocalyx composition using complementary approaches. There are many functions of the BBB glycocalyx; assessing each was outside the scope of this study, so we chose to focus on its barrier function due to its relevance to the BBB. However, future studies should explore the other functions of the BBB glycocalyx in health and disease. Lastly, we decided to focus on EAE as a model for multiple sclerosis, as there was evidence for BBB glycocalyx disruption in this disease and because of the high rates of leukocyte trafficking, a process in which there is evidence that the BBB glycocalyx is intimately involved^19,22,69^. Our findings may differ in other neuroinflammatory conditions or in PV glycocalyces in other locations throughout the body. Nevertheless, some of the methods we put forward may improve our understanding of the BBB glycocalyx in health and disease.

In conclusion, this study provides a detailed characterization of the BBB glycocalyx, highlighting its distinct structure, molecular composition, and barrier function compared to PV glycocalyces. By building on existing techniques, we overcame significant methodological challenges, uncovering the BBB glycocalyx’s resilience during neuroinflammation and its potential role in immune cell trafficking. Targeting the BBB glycocalyx could offer novel therapeutic strategies to modulate immune cell access to the brain while maintaining BBB integrity, opening new avenues for treating neuroinflammatory diseases and other CNS disorders.

## Methods

### Animals

C57BL/6J mice were bought from Jackson Laboratories (strain: #000664) for experiments at the University of California, San Diego (UCSD) or from Janvier Labs (strain: C57BL/6JRj) for experiments performed at the University of Copenhagen (two-photon imaging). All experiments were performed under Institutional Animal Care and Use Committee (IACUC) approval at UCSD or the Federation of European Laboratory Animal Science Associations (FELASA) at the University of Copenhagen. Endothelial HS knockout mice were generated by crossing Extl3^f/f^ (MGI:3844733)^70^ mice with Cdh5-CreER^T2^ mice (MGI:3848982)^71^. Tamoxifen (2 mg in 100 µl corn oil) was administered IP for 3 consecutive days at 5 weeks of age and experiments were performed at 11-13 weeks of age. Brain endothelial sialic acid knockout mice were generated by crossing Cmas^f/f^ (MGI: 6281180) mice with Slco1c1-Cre mice (Mark Kahn, University of Pennsylvania). Mice were housed in a 12 h light/dark cycle with 2-5 mice per cage and were used at 2-4 months of age.

### Reagents

For more detailed information about reagents, see **Supplementary File 5.**

### Electron microscopy

Mice were prepared by a modified version of a previously described protocol^33,34^. Mice were first anesthetized with ketamine (240 mg kg^−1^) / xylazine (36 mg kg^−1^). Mice were then transcardially perfused at 8 ml min^−1^ with a peristaltic perfusion pump for 45 s with mammalian ringer’s buffer^68^ followed by 5 min of lanthanum nitrate fixative (2% lanthanum nitrate, 2% sucrose, 2% glutaraldehyde in 0.1 M sodium cacodylate buffer, pH 7.4). Critically, if the liver did not rapidly clear within ∼20 s and the tail did not vigorously spin within ∼1 min of fixative perfusion, the mouse was abandoned since anything other than ideal perfusion conditions can significantly reduce glycocalyx integrity (data not shown). Fixative was prepared fresh before perfusion and air exposure was limited as lanthanum can precipitate as LaCO_3_ with prolonged storage or extended exposure to air^68^. After perfusion, organs were immediately removed and 1 mm^3^ cubes were cut from the somatosensory cortex (brain), hamstring (muscle), left ventricle (heart), renal cortex (kidney), left lateral lobe (liver), or lumbar spinal cord. Cubes were placed directly into the lanthanum nitrate fixative and stored at room temperature for 4 h and then at 4 °C overnight or up to 1 week before processing at the UCSD CMM Electron Microscopy Core. The cubes were then briefly soaked in a NaOH solution (0.03 M NaOH in 2% sucrose). Note: After the experiments in Fig. 1a-b, cubes were additionally fixed with 1% OsO_4_ in 0.15 M sodium cacodylate buffer for 1-2 h on ice, washed 5 × 10 min in 0.15 M sodium cacodylate buffer and rinsed in ddH_2_O on ice as this improved tissue integrity and staining and did not affect glycocalyx structure. Samples were then incubated in 2% uranyl acetate for 1-2 h at 4 °C, dehydrated at increasing concentrations of ethanol on ice and then dry acetone for 15 min at room temperature, placed in a 50:50 ethanol/Durcupan mixture for 1 h at room temperature and then incubated in 100% Durcupan overnight. Tissue was embedded in Durcupan at 60 °C for 36-48 h. Ultrathin sections (60 nm) were cut on a Leica ultramicrotome with a diamond knife followed by staining with uranyl acetate and Sato lead.

Blood vessels were then imaged with a JEOL 1400 plus. 10-25 circular capillaries per sample (long diameter < 2 × short diameter, diameter < 8 µm, with no significant tears) were quantified with Fiji ImageJ. Total thickness was calculated by taking the average thickness of the glycocalyx across 8 evenly spaced locations around the vessel. Dense thickness was calculated in the same manner but measuring only the portion of the glycocalyx where individual strands were indistinguishable. Glycocalyx area was defined as the percentage of the vessel lumen the glycocalyx occupied as measured using the thresholding function. Coverage was defined as the percentage of the vascular surface where any glycocalyx was present. Arterioles and venules were defined as vessels with a short diameter > 10 µm and arterioles were differentiated from venules based on the presence of block shaped smooth muscle cells lining the vessels as previously described^72^. These vessels were quantified without the requirement of a circular vessel due to their limited number and large size and area was not quantified due to the inconsistent shape of the larger vessels.

### Endothelial transcriptomics

475 glycosylation related genes were identified including glycosyltransferases^36^, glycan degradation enzymes^37,38^, canonical and high-confidence putative mucins^39^, and proteoglycan core protein/synthesis/sulfation genes. From a previous endothelial RNA sequencing study, gene expression in ECs was identified across organs in health and in spinal cord ECs in EAE^40^ **(Supplementary File 1)**. The PCA plot was generated based on expression levels of all 475 genes in ECs from each organ using the prcomp function in R. Glycopacity was used to illustrate BBB specificity of all glycosyltransferases^42^. BBB enrichment and BBB specificity were calculated as previously described^40^: BBB Enrichment = [(log_2_(BE + 0.1) − log_2_(HE +0.1)) + (log_2_(BE + 0.1) − log_2_(KE + 0.1)) + (log_2_(BE + 0.1) − log_2_(LuE + 0.1)) + (log_2_(BE + 0.1) − log_2_(LiE + 0.1))]; BBB Specificity = [BBB Enrichment]^3^ × BE; BE, brain endothelial c.p.m., KE, kidney endothelial c.p.m., LuE, lung endothelial c.p.m., LiE, liver endothelial c.p.m.. To compare genes with proteomics data, mouse genes were assigned from proteins to find the overlap in the three datasets **(Supplementary File 2)**. In EAE, like in the original publication^40^, upregulation was defined as log_2_(fold change) > 1 and expression > 5 c.p.m. in the disease condition, with P < 0.05, and downregulation was defined as log_2_(fold change) < −0.8 and expression of > 5 c.p.m. in the control, with P < 0.05 (N = 3, Wald test). The acute timepoint was labeled onset and subacute was labeled peak for consistency, as they described the same timepoints of the disease model.

### Glycomics

Mice were prepared as described previously with minor changes^44^. Briefly, mice were transcardially perfused with a peristaltic pump for 5 min at 5 ml min^−1^ with ice cold DPBS to remove blood, followed by ice cold 0.5 mg ml^−1^ sulfo-NHS biotin in PBS (prepared fresh) for 10 min at 3 ml min^−1^, followed by ice cold 50 mM Tris-HCl for 5 min at 3 ml min^−1^. The brain, spinal cord, both kidneys, and medial and left lateral lobes of the liver were harvested and homogenized in homogenization buffer with zirconia beads and a BeadBeater. Muscle was not analyzed due to incompatibilities during the homogenization and filtration steps and heart was not analyzed due to concerns of non-glycocalyx labeling throughout the chambers of the heart. The supernatants were isolated, spun through 0.45 µm filters (Corning Spin-X), and stored at −80 °C. On the day of isolation, filtered supernatants were thawed. For each sample, columns with 500 µl settled streptavidin agarose resin were pre-washed 3× with 50 mM ammonium bicarbonate. The samples were then incubated on the columns on a rocker at room temperature for 30 min and washed 5× with ammonium bicarbonate. To elute the samples, the samples were incubated with 10 µg of trypsin in 500 µl ammonium bicarbonate for 2 h at 37 °C and heat-inactivated at 95 °C for 15 min. BCA assays were used to quantify eluted protein concentration, per manufacturer’s instructions, before processing at the UCSD GlycoAnalytics core. From there, the samples were divided and prepared for sialic acid, HS, CS, and N-glycan analyses. For biotin localization staining, mice were perfused for an additional 10 min with 4% PFA at 5 ml min^−1^ followed by staining and imaging as described below.

For hyaluronan glycomics, the samples were prepared the same with a few adjustments. 1 mM EDC was added to the sulfo-NHS biotin solution. 250 µl of streptavidin agarose resin per sample was used. After BCA protein quantification, an additional hyaluronidase treatment step (100 µg in 500 µl ammonium bicarbonate for 1 h at 37 °C) was added to elute any hyaluronan that was still bound to the column.

For sialic acid analysis, the samples were hydrolyzed using 2 M acetic acid at 80 °C for 3 h to release sialic acids. The released sialic acids were purified by spin filtration using a 3,000 MWCO filter and tagged with DMB reagent. The fluorescence-tagged sialic acid derivatives were then analyzed by reverse-phase HPLC with an online fluorescence detector.

To isolate HS and CS, the samples were passed through a DEAE column, and the bound GAGs were eluted with 2 M NaCl. The GAGs were then desalted on PD10-size exclusion column and purified GAGs were lyophilized and used for further analysis. GAGs were then dissolved in HS digestion buffer and reacted with a mixture of Heparinase I, II and III or CS digestion buffer and reacted with a mixture of Chondroitinase ABC at 37 °C for 18 h. Following digestion, the mixtures were fractionated using a 10,000 MWCO filter to remove the enzyme and undigested GAG chains. The supernatant was dried and used for further analysis. For hyaluronan, samples were lyophilized, dissolved in hyaluronan digestion buffer and digested with hyaluronidase at 37 °C for 18 h. Aniline-tagged disaccharides were then separated on a C18 column using an ion pairing solvent mixture and analyzed by mass spectrometry in negative ion mode.

N-glycans were isolated using recombinant PNGase-F. Released N-glycans were applied to tandemly connected Sep-Pak C18 and PGC cartridges equilibrated in water. PGC bound glycans were eluted with 30% acetonitrile containing 0.1% TFA in water, lyophilized and used for further analysis. From there, N-glycans were Per-O-methylated and analyzed by MALDI-TOF/TOF. To do this, samples were dissolved in dry DMSO and stirred for several hours until the samples were completely dissolved. NaOH slurry in DMSO was added followed by the addition of methyl iodide and reacted at room temperature for 45 min. The reaction was stopped by adding 1 ml of ice-cold water. The methylated glycans were extracted using chloroform, dried and used for further analysis. Samples were mixed with super-DHB matrix solution in 1:1 (v/v) ratio and spotted on MALDI plate. The samples were dried and analyzed in positive ion mode. MALDI-TOF/TOF runs we manually annotated and verified using GlycoWorkBench^73^. The proposed assignments for the peaks were based on knowledge of mammalian glycan synthesis pathways.

Glycan quantifications were normalized to total eluted protein following streptavidin isolation as measured by BCA assay. Charge contribution of each glycan was calculated as follows:

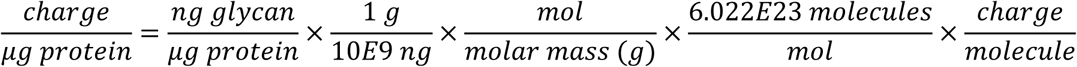

Where 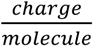 is −1 for sialic acid and hyaluronan and -(1 + average sulfation) for HS and CS. Sialic acid molar mass is 310.76 g mol^−1^, hyaluronan disaccharide molar mass is 379 g mol^−1^ and HS and CS molar mass is calculated based on disaccharide composition/sulfation.

### Glycan staining

To generate tissues for glycan staining across organs, mice were first anesthetized with ketamine (240 mg kg^−1^) / xylazine (36 mg kg^−1^). Then, mice were transcardially perfused with DPBS at 8 ml min^−1^ for 2 min to remove blood. The tissues were then dissected, flash frozen in optimal cutting temperature (OCT) on dry ice and cut into 15 μm sections with a cryostat. For biotin localization staining, organs were incubated in 30% sucrose in PBS overnight at 4 °C, before OCT embedding and sectioning. For semithin sections of individual vessels, endothelial HS knockout and littermate control mice were transcardially perfused with DPBS at 5 ml min^−1^ for 2 min to remove blood and for 10 min with 4% PFA to fix the tissue. 1 mm^3^ cubes were cut from the cerebral cortex and incubated in PBS overnight. 300 nm sections were then cut with a Leica ultramicrotome with a diamond knife.

To stain the tissue, sections were first washed 3× in PBS. If applicable, sections were treated with enzymes as indicated in **(Supplementary File 5)** for 1 h at 37 °C. Sections were then fixed on the slide with 4% PFA in PBS for 10 min and washed again 3× in PBS (samples that were already fixed by perfusion were not fixed again on the slide). When imaging biotinylated probes (FGF2, StcE probe, HABP, MALII) across organs, endogenous biotin signal was blocked with avidin/biotin blocking system per manufacturer’s instructions. Tissue sections were then blocked/permeabilized with 10% goat serum and 0.5% Triton X-100 in antibody buffer. Tissue sections were incubated in primary antibodies overnight at 4 °C and secondary antibodies for 2 h at room temperature with 1% goat serum and 0.05% Triton X-100 in antibody buffer. For HABP staining, tissues were blocked/permeabilized in 2% BSA and 0.5% Triton X-100 in antibody buffer and incubated with antibodies in 0.2% BSA and 0.05% Triton X-100 in antibody buffer. Tissue sections were then dried and coverslipped using Vectashield mounting media with DAPI. Sections were then imaged using a Zeiss Axioimager with Apotome 3 and monochromatic AxioCam 712 with Zen software and quantified with Fiji ImageJ. For glycan staining across organs, images were contrasted to show glycan localization and minimize background, as indicated by no primary control slides.

### Enzyme injections

Under isoflurane anesthesia, mice were given a 100 µl retroorbital injection with enzymes as described in (**Supplementary File 5)** 3 h prior to perfusion, except StcE which was given 24 h before. For the enzyme cocktail group, mice were injected with StcE 24 h before perfusion and injected with the same amount of the remaining enzymes in 100 µl PBS. Timing for enzymatic treatments for two-photon experiments was performed as described in Fig. 4b.

### Primary brain EC culture

Primary brain ECs were isolated from endothelial HS knockout (Extl3^f/f^; Cdh5-CreER^T2^) and littermate controls (Extl3^f/f^) as described previously with slight modificiations^74^. For the first 48 h, cells were cultured with puromycin (4 µg ml^−1^) to kill cells other than brain ECs. Additionally, cells were grown on human collagen type IV (100 µg ml^−1^) and bovine fibronectin (10 µg ml^−1^) coated slide plates. 5-7 days after isolation, monolayers were collected for RT-qPCR or prepared for staining. For immunofluorescence staining, monolayers were fixed with 1% PFA for 10 min, washed with PBS, and stained as described above, with a 30 min block, 1 h primary antibody incubation, and 45 min secondary antibody incubation at room temperature. For RT-qPCR, RNA isolation was performed with the Qiagen RNeasy Micro kit. PrimeTime qPCR primers for Extl3 and β-actin were purchased from Integrated DNA Technologies and cDNA was made using iScript Reverse Transcription Supermix. RT-qPCR was performed using SYBR Green Master Mix. The 2^ΔΔCT^ method normalizing to β-actin was used to calculate relative gene expression of *Extl3*.

### Sodium fluorescein assay

After enzyme injections and 1 h before perfusion, mice were IP injected with 10 µl g^−1^ of 5 mg ml^−1^ NaFl. Just before perfusion, 100 µl of blood was collected via cardiac puncture and stored in an EDTA tube until all perfusions were finished (∼3 h). Mice were then transcardially perfused with PBS at 8 ml min^−1^ with a peristaltic pump for 2 min to remove the blood. Brains were harvested, placed in a tube with PBS (2.5 µl mg^−1^ brain mass) and zirconia beads, and homogenized with a BeadBeater. Brain homogenate and blood samples were then centrifuged, and the brain supernatant and blood plasma were stored at −80 °C until quantification. To quantify samples, plasma (diluted 1:200 in PBS) and brain homogenates were diluted 1:1 in TCA buffer to precipitate proteins at 4 °C overnight, centrifuged, and supernatant was isolated. Supernatants were diluted 1:1 in borate buffer and fluorescence intensities were measured on a plate reader (Tecan Infinite M Plex, excitation 480 nm, emission 538 nm). After background subtracting uninjected control brain and blood values, the brain:blood fluorescence intensity ratios for each sample were calculated.

### Two-photon imaging

The surgical preparation was performed as previously described ^24,75^. Briefly, mice were anesthetized using IP bolus injections of xylazine (10 mg kg^−1^) and ketamine (60 mg kg^−1^) to induce sedation. Subsequently, anesthesia was maintained with additional IP injections of ketamine (30 mg kg^−1^) administered every ∼20 minutes. A tracheotomy was performed to enable mechanical ventilation. Ventilation was maintained using a MiniVent Type 845 respirator (Harvard Apparatus) with a tidal volume of 180–220 µl and a respiration rate of 190–240 strokes min^−1^. The inhaling air was supplemented with oxygen at a flow rate of 1.5–2 ml min^−1^ to ensure adequate oxygenation. Continuous real-time monitoring of end-tidal CO_2_ levels (Capnograph Type 340, Harvard Apparatus) was employed to maintain physiological conditions, with acceptable ranges of exhaled CO_2_ maintained at 2.2–2.8%. Dual catheterization was carried out by inserting one catheter into the femoral artery for real-time monitoring of mean arterial blood pressure (50–80 mmHg, monitored via BP-1 Pressure Monitor, World Precision Instruments) and delivery of fluorescent dyes. Another catheter was inserted into the femoral vein for the administration of anesthesia during imaging. This setup ensured precise control over systemic physiological parameters.

To allow for optimal two-photon imaging, a cranial window was surgically prepared over the somatosensory barrel cortex. Right before cranial window surgery, enzymes were injected retroorbitally except StcE, which was injected 21 h before the procedure. After securing the skull with a custom metal head plate using Loctite superglue, a 3-4 mm craniotomy was performed using a diamond-coated dental drill operating at 7000 rpm. To minimize heat-induced tissue damage, the skull was regularly irrigated with room-temperature saline during the drilling process. Following the removal of the bone flap, the dura mater was carefully excised to expose the cortical surface. To stabilize and protect the exposed brain tissue, a drop of 0.75% low-melting-point agarose dissolved in artificial cerebrospinal fluid was applied. The craniotomy was then sealed with a glass coverslip of 0.08 mm thickness, ensuring optical clarity for imaging. Next, the animal was transferred to the imaging stage, and the anesthesia was switched to α-chloralose (50 µg h^−1^ g^−1^) infused via a venous catheter. During all steps of imaging, the exhaled CO_2_ and blood pressure were continuously monitored.

The imaging was performed with an SP5 upright laser scanning two-photon microscope (Leica Microsystems) equipped with a MaiTai Ti:Sapphire tunable infrared laser (Spectra-Physics), using 20×, 1.0 NA water-immersion objective with a planar optical resolution of approximately 0.53 µm in the XY plane and 2.5 µm in depth^76^. Fluorescence signals were collected using Leica Advanced Fluorescence software (version 2.7.3.9723) by two separate multi-alkali detectors after passing through the dichroic mirror and appropriate bandpass filters (525–560 nm for green-emitting fluorophores, and 560–625 nm signals from red-emitting fluorophores).

After being mounted for imaging, a bolus injection of 2 µl g^−1^ 1% Alexa 488 conjugated albumin was given through the arterial catheter. A single field of view (0-150 µm depth) was imaged right before albumin injection, right after, and then once every five minutes for 1.25 h at 900 nm excitation (540 mW at the objective). Data were collected as 16-bit image hyperstacks (Z-stacks over time) at 512 x 512 pixel resolution (775 µm x 775 µm, Z-step = 5 µm). Then, 300 µl 0.5 mg ml^−1^ Texas Red conjugated Lycopersicon Esculentum lectin was administered through the arterial catheter with a syringe pump over 5 min (3.6 ml h^−1^). About 3 h after albumin injection, the laser power was decreased (65 mW at the objective), which minimized albumin fluorescence. Next, a bolus injection of 2 µl g^−1^ 1% NaFl was given through the arterial catheter. One field of view (0-150 µm depth) was imaged right before NaFl injection, right after, and once per minute thereafter for 1 h at 900 nm excitation. Data were collected as 16-bit image hyperstacks at 1024 × 1024 pixel resolution (775 µm x 775 µm, Z-step = 5 µm). Hyperstack images were then analyzed using Fiji ImageJ. Hyperstacks were first stabilized using the plugin image stabilizer^77^ and Z-projected using average signal intensity projection from the top of the brain to 100 µm into the parenchyma (20 Z-planes), and plugin image stabilizer^77^. The parenchymal intensity was then calculated by measuring the average intensity over time in 100 circular regions of interest (ROI) where there was no vasculature present. To calculate plasma intensity, hyperstacks were Z-projected across the same 100 µm depth using maximal signal intensity projection, and the average signal was measured in 20 circular ROI within the vasculature. Background intensities, calculated from the same ROI before injection of albumin or NaFl, were subtracted from the parenchymal and plasma intensities, and then parenchymal intensities were divided by plasma intensities to get brain:blood fluorescence intensity ratios.

### Experimental autoimmune encephalomyelitis

EAE was induced using a commercial kit (Hooke Laboratories, EK-2110) as described previously^78^, with a slightly modified scoring protocol. Briefly, after 7 days of acclimatization, mice were given two subcutaneous MOG/CFA injections on the upper and lower back and one intraperitoneal pertussis toxin injection (110 ng) under isoflurane anesthesia on day 0. On day 1, mice were given a single pertussis toxin IP injection under isoflurane anesthesia. Mice were weighed and scored daily as follows:

- 0.5 = Tip of tail is limp
- 1.0 = Entire tail is limp
- 1.5 = One hind leg falls through wire rack consistently when walking
- 2.0 = Legs and toes do not spread when lifted or one foot is dragging
- 2.5 = Both hind feet are dragging but legs have movement, or one leg is not being used to walk
- 3.0 = Both hind legs are not being used to walk but legs can move
- 3.5 = Both hind legs are immobile or mouse is unable to right itself when placed on its side
- 4.0 = Partial front leg paralysis, but mouse can move (if 4.0 for 2 straight days, euthanize)
- 4.5 = Partial front leg paralysis, mouse cannot move (euthanize)
- 5.0 = Death/euthanasia/spontaneous rolling (euthanize)

The onset timepoint was defined as the first day of scoring or first date of weight loss greater than 1 g over a 2-day period. The peak timepoint was defined as 2-5 days after onset with a score ≥ 2.5 and all peak EAE mice for the glycomics experiments were perfused on day 17 with the same requirements. The chronic timepoint was defined as day 21-30 with a max score ≥ 2.5. After onset of hind leg paralysis, mice were given access to wet food.

### Statistics

The statistics for each experiment are described in the figure legends. All statistics were calculated using GraphPad Prism software (version 10). In the text, all ± values indicate standard deviation. In figures, all error bars represent the standard error of the mean, and N values refer to the number of animals, or pooled replicates for primary EC and glycomics experiments. All graphs were made using GraphPad Prism software (version 10), except Fig. 2a-b and Extended Data Fig. 1 which were made in R (version 4.1.3). Sample sizes were selected based on the variability of the measurement and values of difference between conditions.

## Supporting information

Supplementary File 1 - Glycosylation Genes Full Data

Supplementary File 2 - Glycosylation Genes with Toledo et al and Tremblay et al

Supplementary File 3 - Albumin accumulation

Supplementary File 4 - NaFl Accumulation

Supplementary File 5 - Reagents

## Acknowledgements

The authors would like to thank the University of California, San Diego - Cellular and Molecular Medicine Electron Microscopy Core (UCSD-CMM-EM Core, RRID:SCR_022039) for equipment access and technical assistance. We also thank Mark Kahn at the University of Pennsylvania for generously sharing the Slco1c1-Cre mouse line. R.L. was supported in part by the UCSD Graduate Training Program in Cellular and Molecular Pharmacology through an institutional training grant from the National Institute of General Medicine Sciences, T32 GM007752 and by the UC San Diego Merkin Graduate Fellows Program. K.K. and M.La. were supported by a grant from the Lundbeck Foundation (#R433-2023-1150 and #R453-2024-359). R.D. was supported by a grant from the Cure Alzheimer’s Fund and NIH R01 NS091281.

## Author Contributions

R. L., J.D.E., and R.D. conceived the project, designed experiments, and wrote the manuscript with input from all authors. R. L. performed the experiments. K.K., and M.La. aided in the design of two-photon experiments and their analysis. K.K. and M.Lø. provided technical guidance and performed surgeries for two-photon experiments. S.A. performed primary brain EC culture experiments. M.K.B.M. performed histological stainings of glycans across organs and analyzed the images. B.C. and M.P. acquired mass-spectrometry and liquid chromatography based glycomic data and aided in data analysis. D.C.L. helped write and edit the manuscript. M.A. and A.M-K. provided the Cmas floxed mice. A.G.T. aided in the design of the glycomics experiments.

---

**Extended Data Figure 1:**
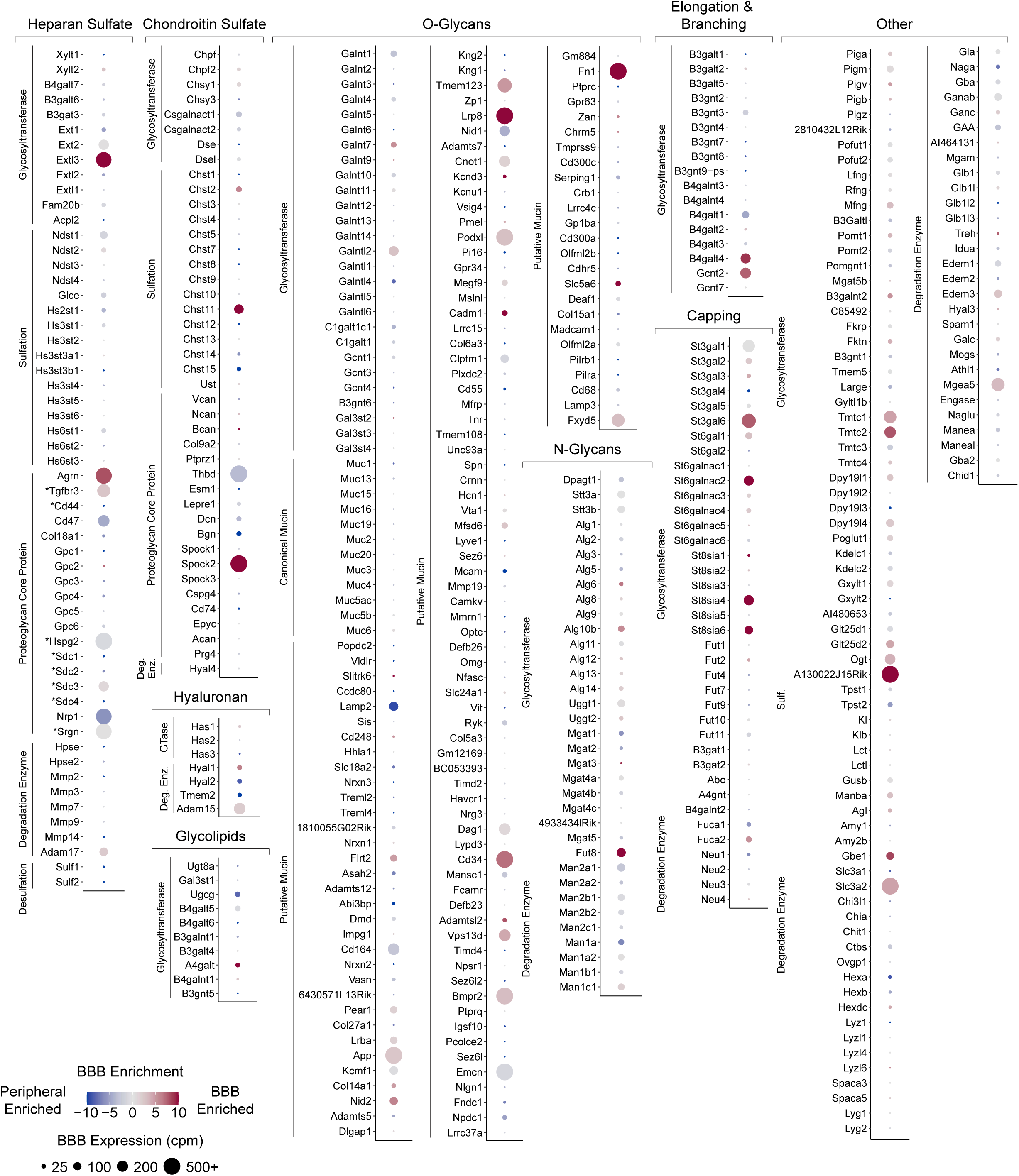
BBB expression and enrichment of glycosylation related genes. BBB expression and BBB enrichment levels of all 475 glycosylation related genes organized by glycan and process. BBB expression level is shown by size of the circle and BBB enrichment is shown by color, with red indicating BBB enrichment and blue indicating peripheral enrichment. * indicates HS proteoglycans that are also known to also contain CS chains. Original data from Munji and Suong et al^40^.

**Extended Data Figure 2:**
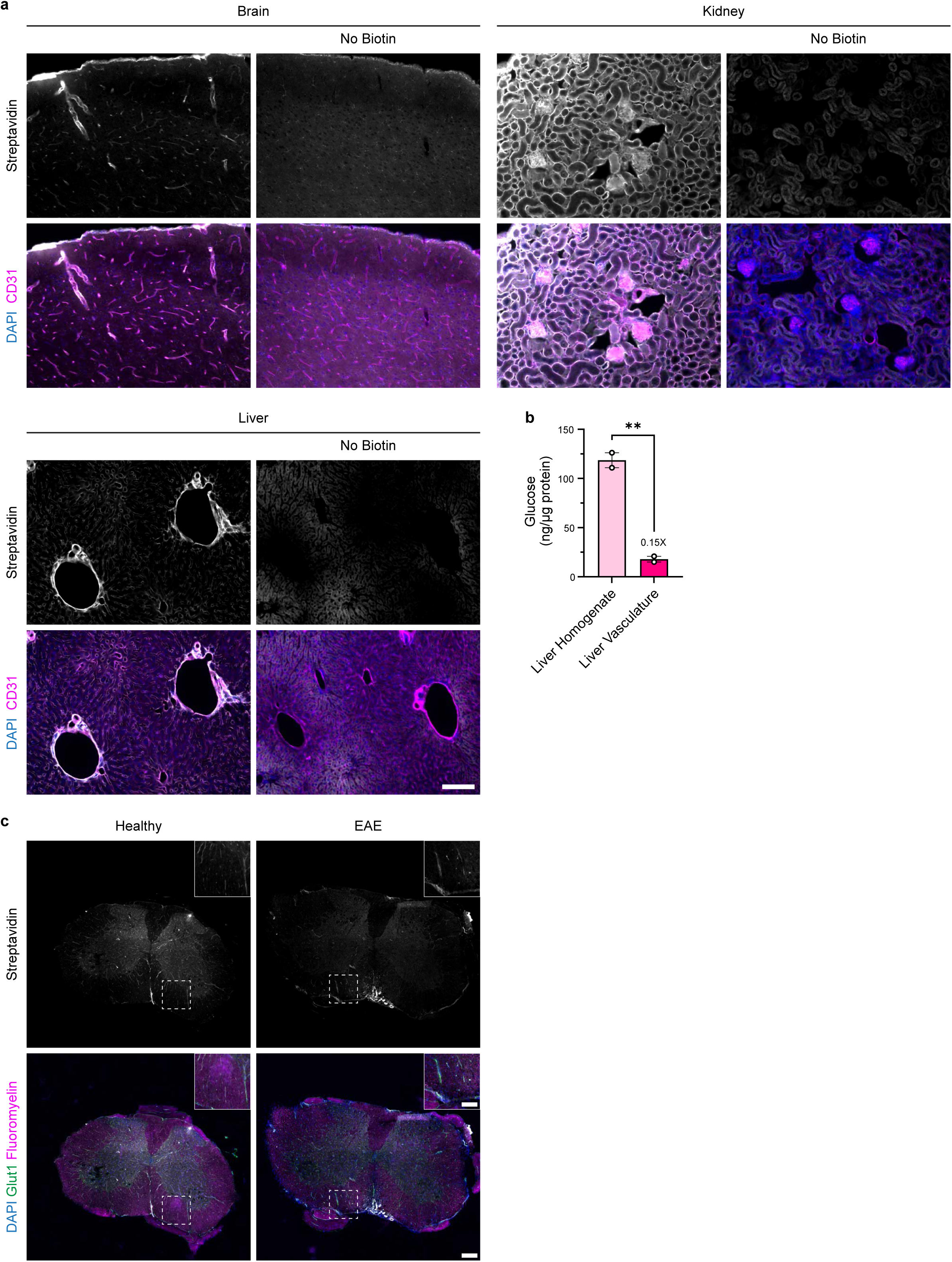
Confirmation of vascular tagging. **a,** Streptavidin staining to show localization of biotin following biotin perfusion for glycomics. Anti-CD31 stains ECs (scale bar = 100 µm). **b,** Relative abundance of glucose in liver homogenate and liver vasculature (N = 2, **P < 0.01, two-sided t-test). **c,** Streptavidin staining to show biotin localization in healthy and EAE lumbar spinal cord sections following biotin perfusions for glycomics. Fluoromyelin stains myelin, and anti-Glut1 stains brain and spinal cord ECs (Full image scale bar = 200 µm, inset scale bar = 100 µm).

**Extended Data Figure 3:**
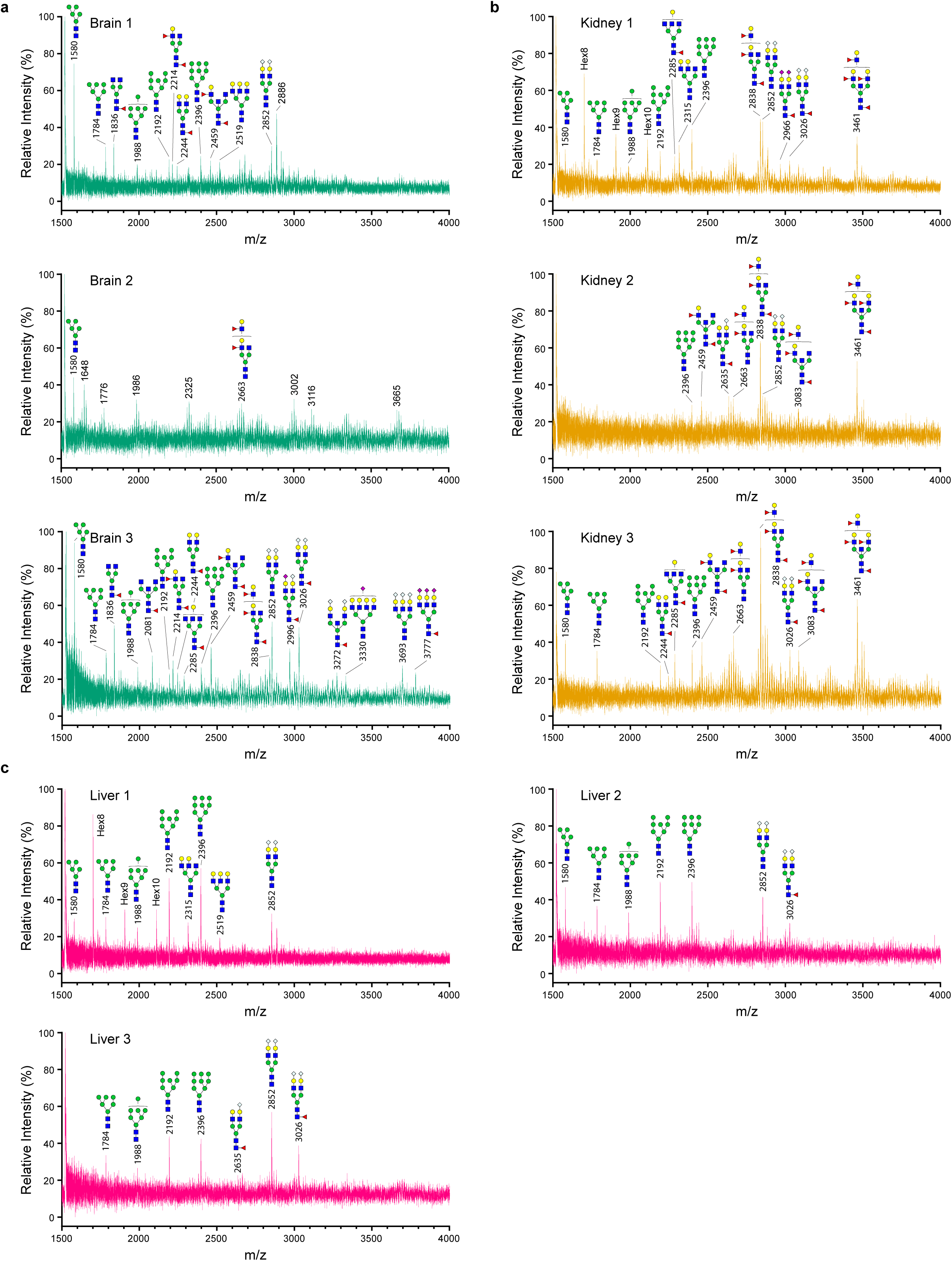
Glycocalyx N-glycans. Annotated N-glycan MALDI-TOF/TOF profiles from biotin tagged **a** BBB, **b** kidney, and **c** liver glycocalyx samples.

**Extended Data Figure 4:**
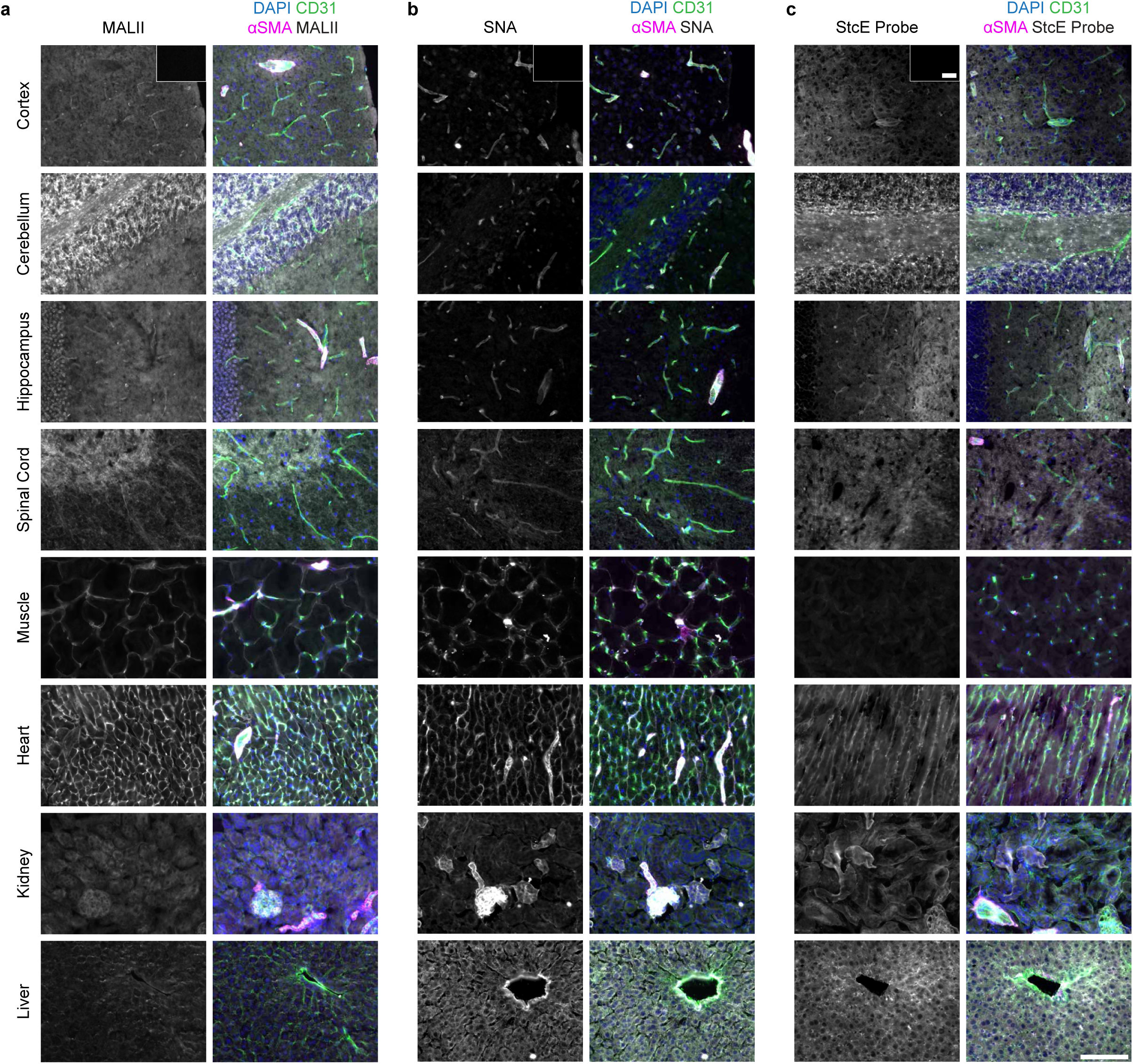
Sialic acid and mucin localization across organs. a-c, Tissue sections from the brain (cortex, cerebellum, and hippocampus), spinal cord, muscle, heart, kidney, and liver were stained with DAPI, anti-CD31 (blood vessels), anti-αSMA (large blood vessels), and the indicated glycan binding protein. **a,** MALII stains α2-3 linked sialic acid. **b,** SNA stains α2-6 linked sialic acid. **c,** StcE probe stains mucins. Cortex inset shows cortex staining following on-slide enzymatic treatment with sialidase **(a-b)** or mucinase StcE **(c)**. Full image scale bar = 100 µm. Inset scale bar = 100 µm.

**Extended Data Figure 5:**
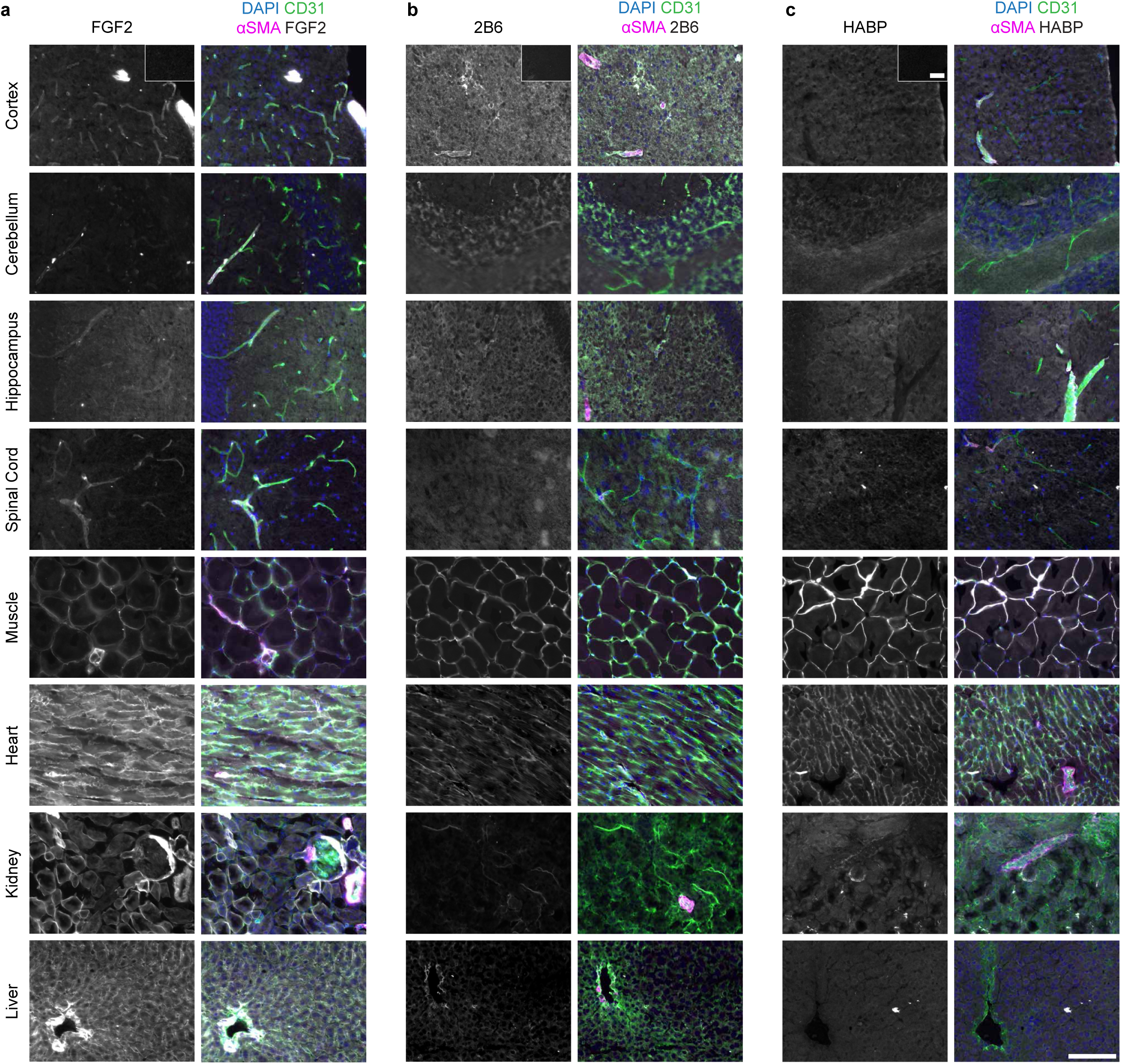
GAG localization across organs. a-c, Tissue sections from the brain (cortex, cerebellum, and hippocampus), spinal cord, muscle, heart, kidney, and liver were stained with DAPI, anti-CD31 (blood vessels), anti-αSMA (large blood vessels), and the indicated glycan binding protein. **a,** FGF2 stains HS. **b,** Slides were stained following enzymatic treatment with chondroitinase ABC to expose C-4-S stubs. 2B6 stains C-4-S stubs. **c,** Hyaluronan binding protein (HABP) stains hyaluronan. Cortex inset shows cortex staining following on-slide enzymatic treatment with heparinase II **(a)**, hyaluronidase **(c)**, or PBS treatment **(b)**. Full image scale bar = 100 µm. Inset scale bar = 100 µm.

**Extended Data Figure 6:**
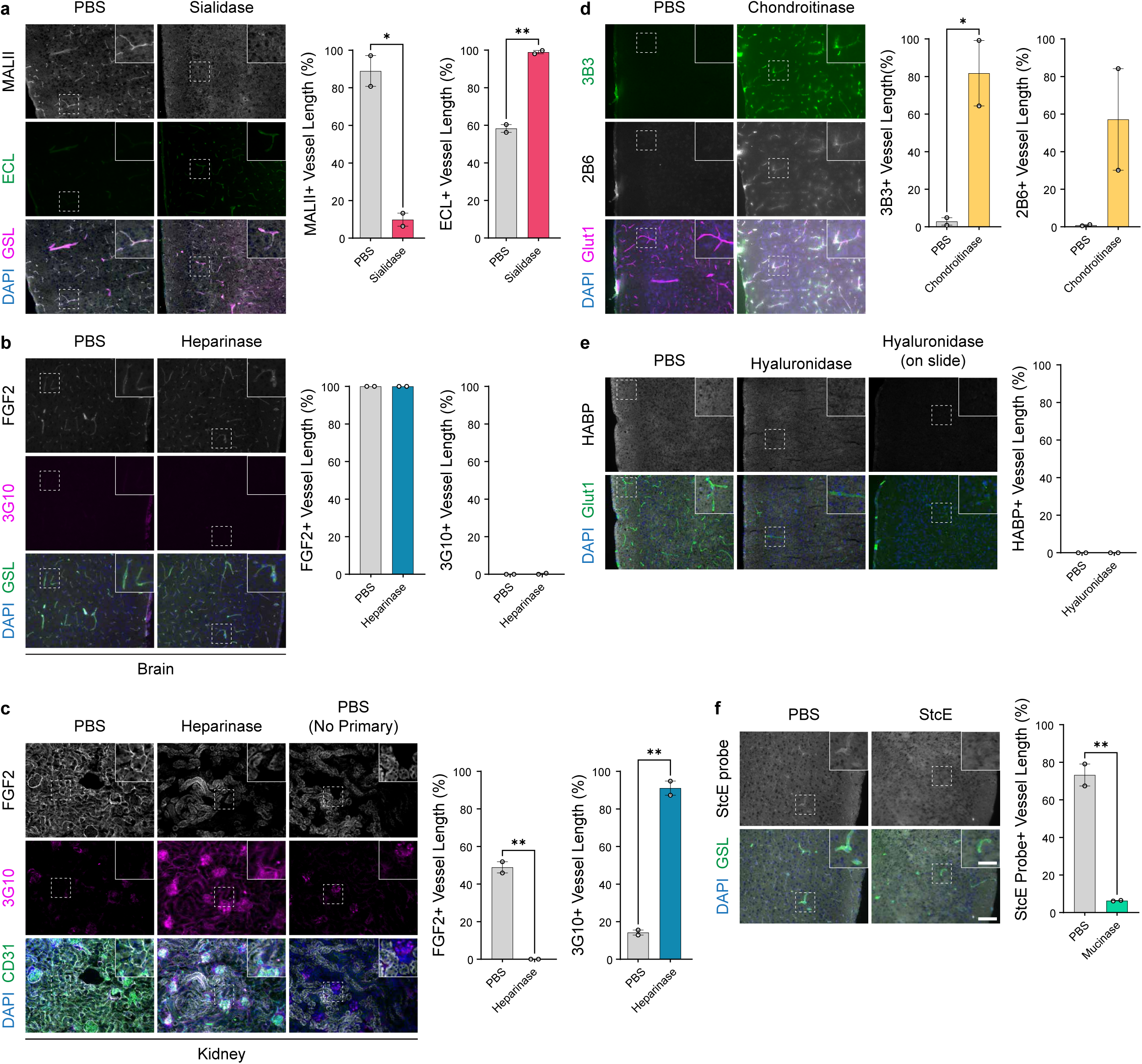
Degradation of BBB glycocalyx glycans by enzymatic injection. a-h. Tissue sections from the cerebral cortex **(a-b,d-f)** or kidney **(c)** following retroorbital injection of PBS or the indicated enzymes. **a,** Staining with MALII (α2-3 sialic acid), ECL (exposed N-acetyllactosamine (LacNAc)) and GSL (blood vessels) and quantification of vascular sialic acid and exposed LacNAc following injection of PBS or sialidase. **b-c,** Staining with FGF2 (HS), 3G10 (HS stub), and anti-CD31 or GSL (blood vessels) and quantification of vascular HS and HS stub following injection of PBS or heparinase I, II, III. **d,** Staining with 3B3 (C-6-S CS stub), 2B6 (C-4-S CS stub), and anti-Glut1 (blood vessels) and quantification of C-6-S and C-4-S stubs following injection of chondroitinase ABC. **e,** Staining with HABP (hyaluronan), and anti-Glut1 (blood vessels) and quantification of vascular hyaluronan following injection of PBS or hyaluronidase. **f,** Staining with StcE probe (mucins) and GSL (blood vessels) and quantification of vascular mucins following injection of PBS or mucinase StcE. Full image scale bar = 100 µm. Inset scale bar = 50 µm (N = 2, mean ± s.e.m., *P < 0.05, **P < 0.01, two-sided t-test).

**Extended Data Figure 7:**
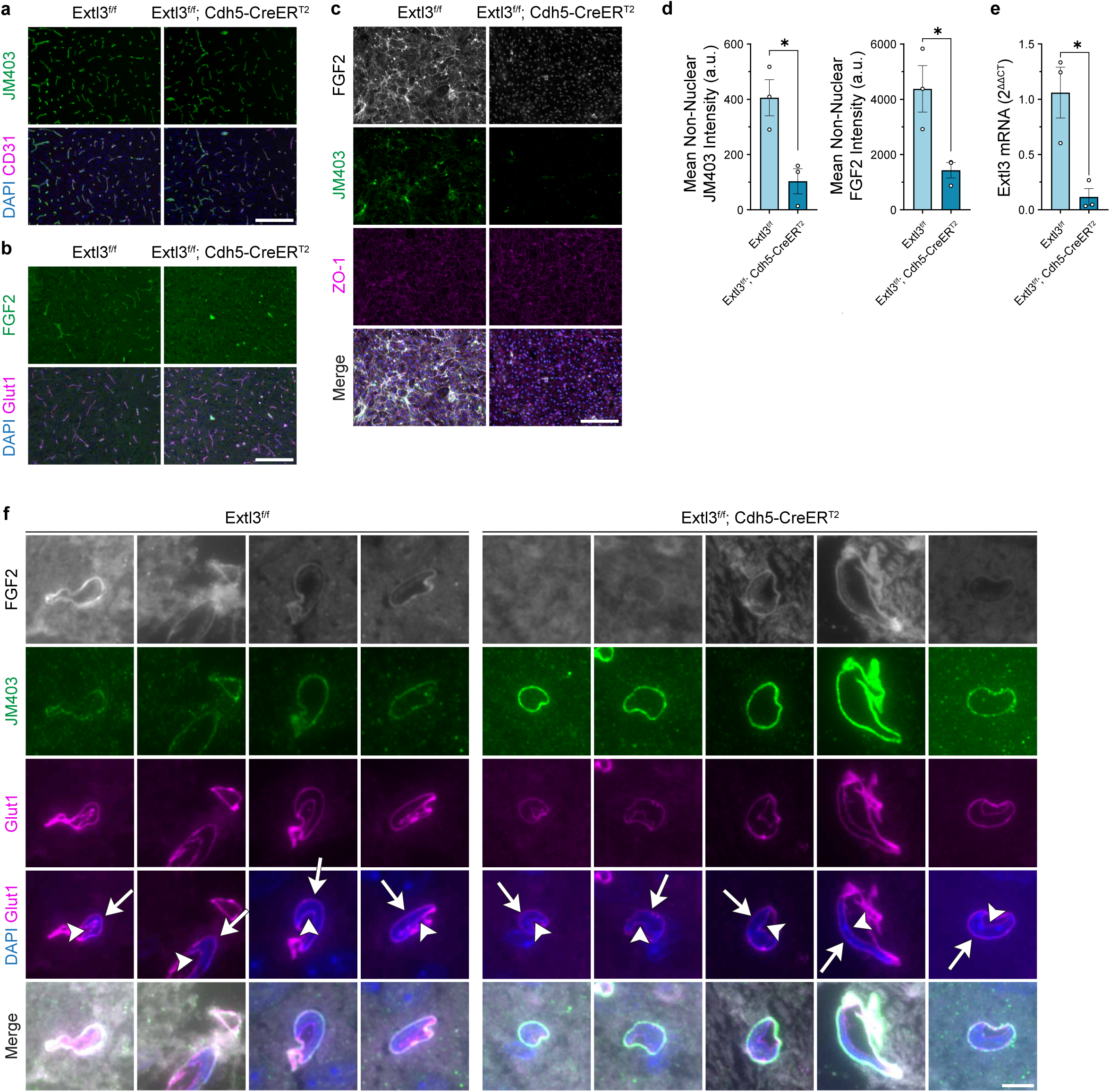
Brain endothelial HS is mostly in the basement membrane. a-b, Staining of the cerebral cortex in endothelial HS knockout mice (Extl3^f/f^; Cdh5-CreER^T2^) or littermate controls (Extl3^f/f^) 6-8 weeks after tamoxifen injection. Staining with JM403, FGF2 (HS), anti-CD31, and anti-Glut1 (blood vessels). Scale bar = 200 µm. **c,** Staining for HS and ZO-1 (blood vessels) in primary brain ECs isolated from littermate control and endothelial HS knockout mice and **d** quantification of non-nuclear HS intensity (N = 3, mean ± s.e.m., *P < 0.05, two-sided t-test). Scale bar = 200 µm. **e,** Quantification of *Extl3* mRNA isolated from primary ECs in littermate controls and endothelial HS knockout mice (N = 3, mean ± s.e.m., *P < 0.05, two-sided t-test). **f,** Staining of individual cerebral cortex capillaries from littermate control and endothelial HS knockout mice from thin (300 nm) sections. Anti-Glut1 stains both the luminal (white arrowhead) and abluminal (white arrow) side of brain ECs. Scale bar = 5 µm.

**Extended Data Figure 8:**
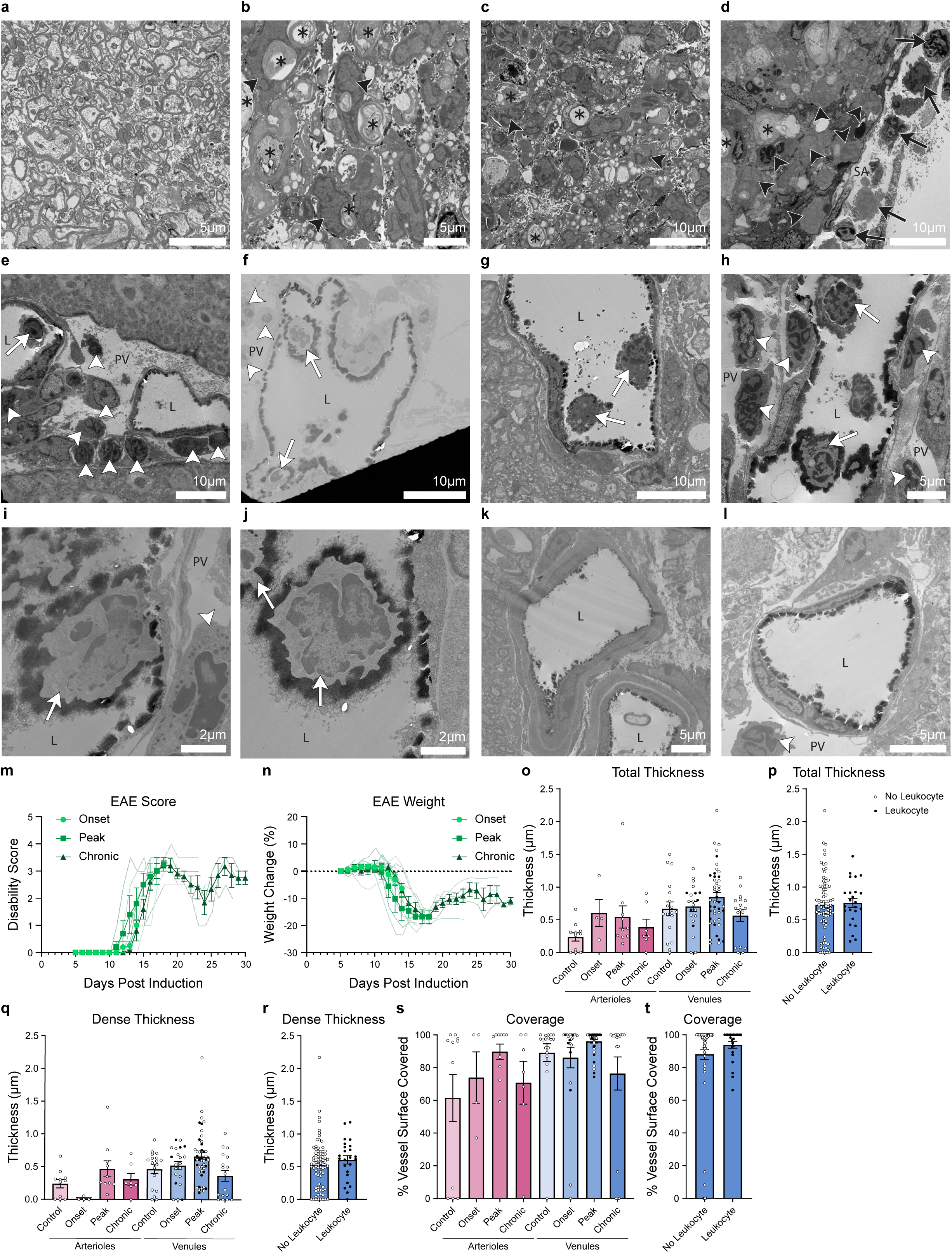
BBB glycocalyx structure during leukocyte extravasation in EAE. EM images of lumbar spinal cord white matter axons in **a** uninjected healthy mice and **b-d** EAE mice. **e-j,** EM images from EAE mice of venules with leukocytes present within the vessel lumen. EM images from EAE mice of **k** arterioles and a **l** venule with no leukocyte in the vessel lumen. Asterisks: dystrophic axons, black arrowheads: leukocytes in parenchyma, black arrows: leukocytes in subarachnoid space, white arrowheads: leukocytes in perivascular space, white arrows: leukocytes in vessel lumen, SA: subarachnoid space, L: vessel lumen, PV: perivascular space. **m,** Disability score and **n** relative weight change of EAE mice taken at each timepoint for EM (N = 5, mean ± s.e.m.). **o-t** Quantification of BBB glycocalyx structural parameters in arterioles and venules of healthy control or EAE mice at each key timepoint of the disease model including **o,p** total thickness, **q,r** dense thickness, and **s,t** coverage. Each dot represents a vessel. **p,r,t** Quantification of venules from all timepoints, including control, with and without a leukocyte present.

**Extended Data Figure 9:**
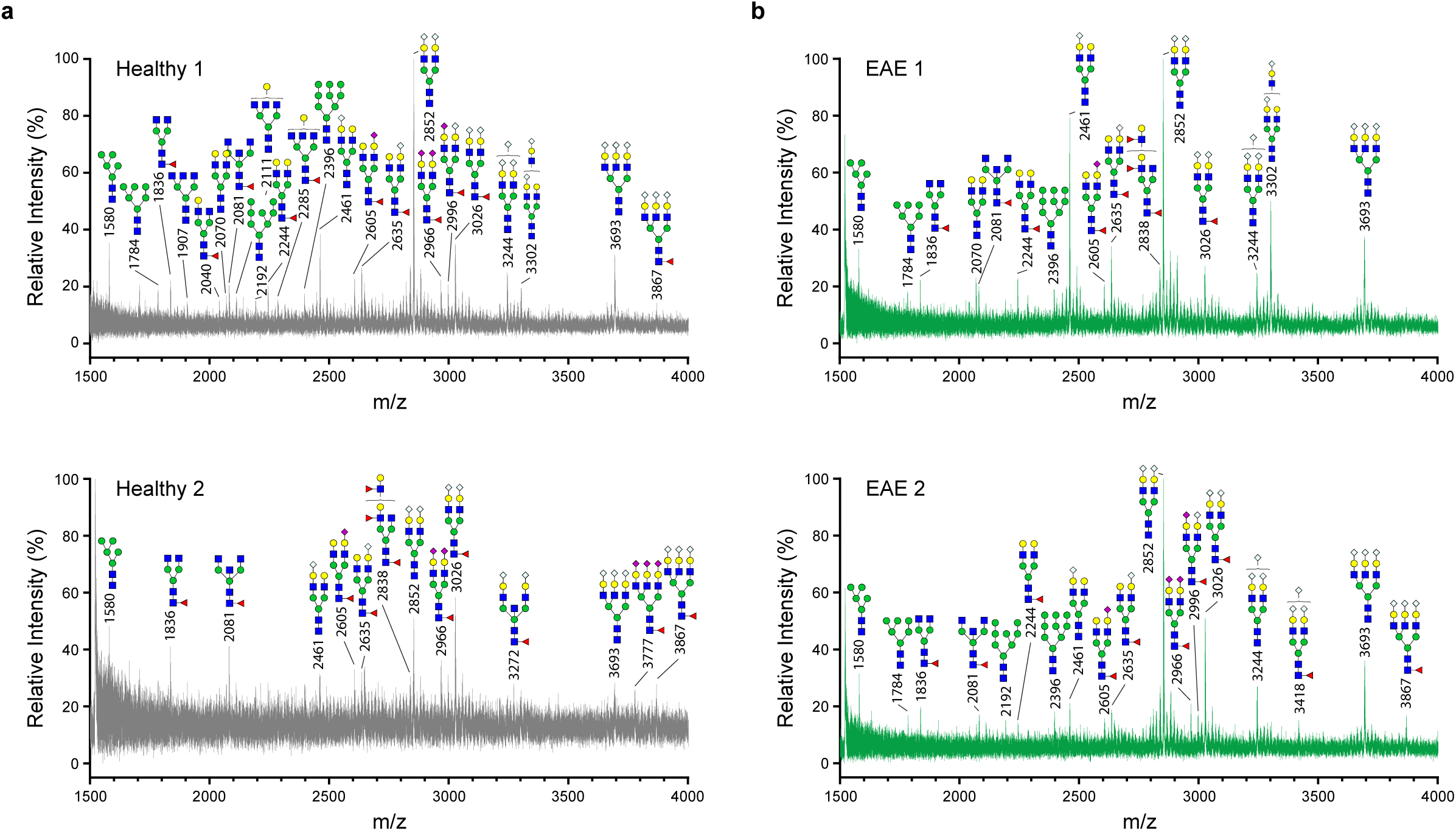
BBB glycocalyx N-glycans in EAE. Annotated N-glycan MALDI-TOF/TOF profiles from biotin tagged spinal cord glycocalyx samples in **a** healthy mice and **b** mice at peak EAE.

## Notes

### Competing Interest Statement

The authors have declared no competing interest.

